# Long-term high temperatures affect seed maturation and seed coat integrity in *Brassica napus*

**DOI:** 10.1101/2024.11.27.625589

**Authors:** Unnikannan Prabhullachandran, Ivana Urbánková, Alejandro Medaglia-Mata, Audrey Creff, Aline Voxeur, Vilma Buršíková, Benoit Landrein, Jan Hejátko, Hélène S. Robert

## Abstract

Temperatures above the optimum growth threshold affect seed development, accelerating embryo development and producing seeds with ruptured seed coats. However, the underlying mechanisms of this effect remain unclear. To investigate temperature-induced seed coat rupture, we used a multidisciplinary approach combining detailed phenotyping, transcriptomics, histology, immunolabelling, cell wall profiling and nanoindentation-based mechanics of the seed coat in oilseed rape *(Brassica napus*). Our data reveal that high temperatures accelerate embryo growth, resulting in larger embryos without a compensatory increase in overall seed size. This rapid embryo expansion may exert elevated mechanical stress on the seed coat cells, significantly thinning the seed coat layers. Concurrently, prolonged exposure to high temperatures drives premature biochemical maturation of the seed coat, characterized by accumulation of demethylesterified pectin. Nanoindentation analysis demonstrated that these changes may have compromised the structural integrity of seed coat cell walls. Ultimately, the weakened seed coat may not withstand the internal tension imposed by the fast-growing embryo, leading to seed coat rupture and reduced seed quality. Our work on seed maturation and the mechanism of heat-induced seed coat rupture provides valuable insights for future research into breeding thermotolerant *Brassica napus*, crucial in the context of a changing climate.

**HIGHLIGHTS:**

- Growth of *Brassica napus* under long-term high temperatures accelerated embryo growth and led to seed coat rupture in approximately half of the seeds.
- Pathways linked to cell wall modification were found to be significantly differentially regulated at high temperatures, accompanied by increased pectin demethylesterification in the seed coat.
- A lower reduced modulus and hardness indicate that the surface of HT seeds is mechanically more compliant and softer. Together with the reduced thickness of the seed coat layers, such a seed coat, as a whole structure, is mechanically weaker and less able to resist deformation imposed by embryo growth, resulting in seed coat rupture.

## 1. INTRODUCTION

The adverse impacts of global warming on agricultural production pose a major challenge to global food security. Extreme weather events, such as heat stress, drought and floods, have the potential to affect the production of staple crops, such as rice, wheat, maize, and soybean (Zhao et al. 2017; Agnolucci and De Lipsis 2020; Jägermeyr et al. 2021; Heino et al. 2023). In 2025, the average annual global temperature was 1.19°C above the baseline average of the 30-year period 1951 to 1980 (Lenssen et al. 2024; GITSTEMP v4 2026), highlighting the need to adapt agricultural practices to higher temperatures.

*Brassica napus* (canola or oilseed rape), an allotetraploid species of the *Brassicaceae* family, is the second most widely cultivated oilseed crop, used for the production of vegetable oil, biofuel, and protein in animal feed. Cultivation of oilseed rape at temperatures above the optimum growth temperature of 21°C for prolonged periods, especially during the flowering stage, has been reported to result in several adverse effects, including rapid vegetative growth, reduced viability of female gametophytes, increased seed abortion rate, accelerated embryo development, and a reduction in seed oil composition (Young et al. 2004; Mácová et al. 2022; Secchi et al. 2023). One of the distinctive phenotypes observed during seed development of certain canola cultivars subjected to prolonged heat stress that affects seed yield is the occurrence of preharvest sprouting of mature seeds prior to their desiccation (Mácová et al., 2022). Preharvest sprouting is a precocious germination of the seeds in the fruit. It has been reported to be the consequence of a hormonal imbalance and absence of seed dormancy (Bewley et al. 2013; Nautiyal et al. 2023). Seed germination, either precocious or after harvest, is associated with the rupture of the seed coat and the protrusion of the embryo radicle. We observed similar phenotypes during seed maturation, referred to, in this study, as seed coat rupture (SCR), and ranging from a mild crack in the seed coat to fully ruptured seeds with protruded embryos (Fig. 1). The exact mechanism by which this phenotype is induced at elevated temperature remains unclear in *Brassica napus*. In *Brassicaceae*, seed size is regulated by the interaction between maternal and zygotic signals that coordinate the proper development of the seed coat, endosperm, and embryo (Li et al. 2019). The interaction between these three genetically distinct seed tissues has been studied in *Arabidopsis thaliana. Arabidopsis* is a widely used model plant, largely due to its extensive availability of genetic resource and its suitability for research using high-resolution microscopy. During the development of *Arabidopsis* seeds, the pressure generated by the interaction between the developing embryo and the endosperm within the seed exerts tension on the developing integuments (Creff et al. 2015, 2023). The generated tension is sensed by a specific layer of the seed coat’s outer integument (oi), the adaxial epidermis (oi1). This mechanical stimulus triggers a transcriptional response marked by the activation of the mechanosensitive gene *EUI-like P450A1* (*ELA1)*, which is believed to promote cell wall stiffening through accumulation of demethylesterified pectin, thereby regulating seed size (Creff et al. 2023). Plant cell expansion depends on cell wall stretching, which is determined by the wall’s mechanical strength and dynamic extensibility (Geitmann and Ortega 2009). The strength and integrity of the wall during growth is maintained actively by the modification of the cell wall elements (Cosgrove 2024). The dry mass of the primary plant cell wall is composed of cellulose, hemicellulose, and pectin (Ayala Soto and Serna Saldívar 2020; Cosgrove 2024). Pectin forms a hydrated gel-like matrix in which cellulose and hemicellulose cross-link to form the cell wall. It is abundant in the middle lamella, an adhesive layer between adjacent cells. The most abundant pectin, homogalacturonan (HG), is produced in a highly methylesterified state. It can undergo demethylesterification by the activity of pectin methylesterase (PME) depending on developmental cues (Turbant et al. 2016; Wu et al. 2018; Xu et al. 2022). Blockwise removal of methyl groups can lead to the formation of Ca^2+^ cross-links, which makes the cell wall more rigid and inhibits growth. Alternatively, random removal can lead to pectin degradation and altered wall rheology (Bidhendi and Geitmann 2016; Wu et al. 2018). Seed size correlates with endosperm proliferation and development. The two processes are linked by a brassinosteroid-induced cell expansion and increased levels of methylesterified pectin in the seed coat, which helps to regulate seed size (Lima et al. 2024).

**Fig. 1.**
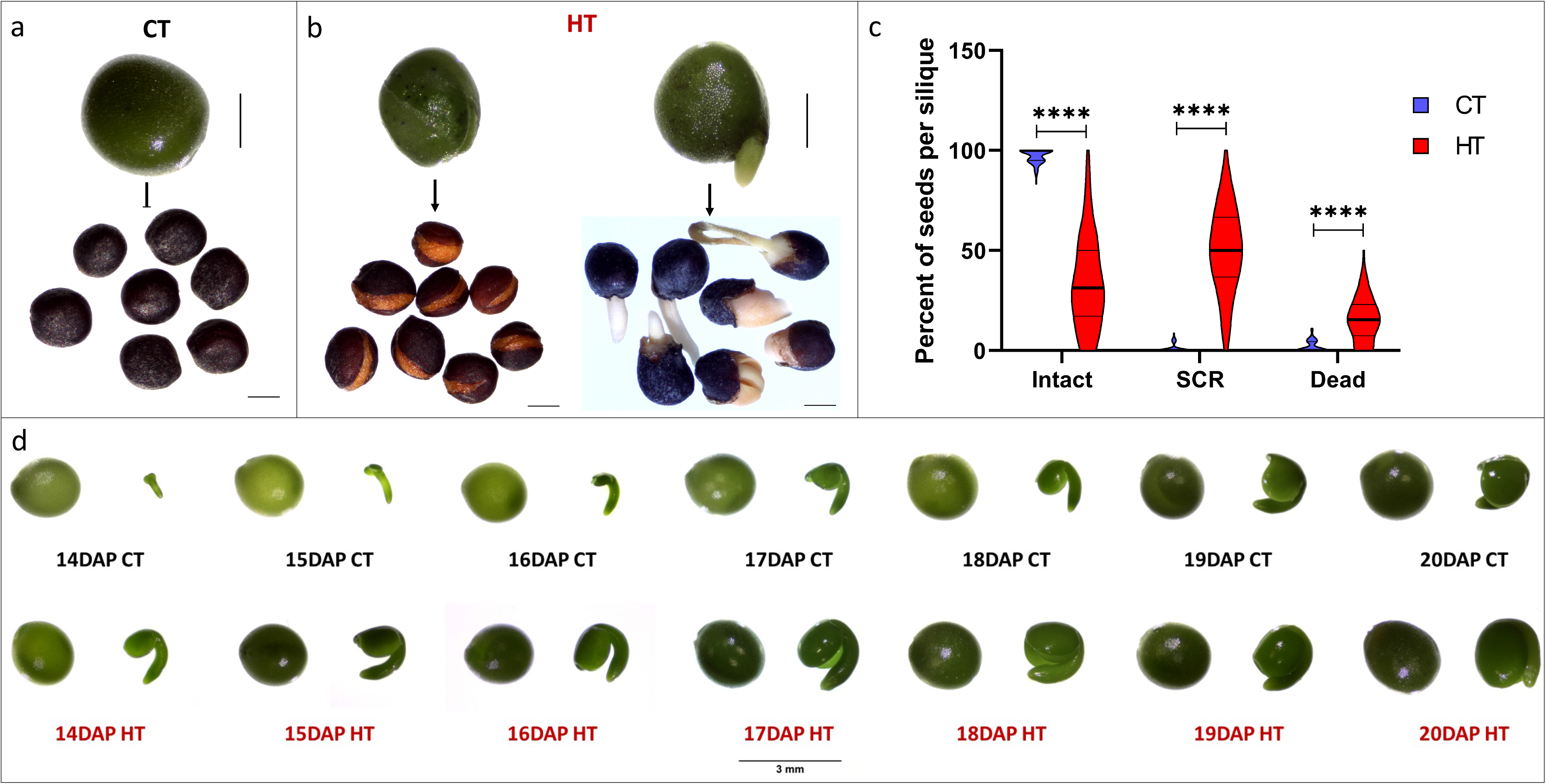
SCR phenotype observed in Topas seeds developed under HT. (a) 26 DAP seeds (top) and matured dry seeds after harvest (bottom) grown under CT. (b) 26 DAP seeds (top) and matured dry seeds after harvest (bottom) grown under HT depicting the seed coat rupture (SCR) with seeds with incomplete seed coat rupture (left) and seeds with fully ruptured seed coat with protruding embryos (right). (c) Percentage of dry seeds per silique from CT (blue) and HT (red) being intact, with SCR or dead (n CT = 116 siliques, n HT = 245 siliques). Phenotypic classification presented in Fig. S1a-c. Asterisks indicate statistically significant differences in HT (**** corresponds to *p*-values of *p* < 0.00005). (d) Embryo stage of seeds developed under CT (top) and HT (bottom) from 14 to 20 DAP. The isolated embryo shown on the right is from the seed on the left. Scale bars: 1 mm (a,b) and 3 mm (d).

In this study, we used a multi-methodology approach to investigate the occurrence of the temperature-induced SCR phenotype in *Brassica napus*. We observed that SCR occurs in seeds around 20 days after pollination (20 DAP) when plants are cultivated at elevated temperatures for a prolonged period. We hypothesized that the pressure induced by unrestricted embryo growth may lead to increased tension in the seed coat, based on our observation of a thinning of the seed coat layers and changes in cell wall composition of the seed coat cells that may have rendered the seed coat structurally weaker. This interpretation is supported by the observation that seeds cultivated at elevated temperatures have a lower reduced modulus and lower hardness in HT seed surface, which may render the seed coat more prone to rupture. The precise mechanism by which heat stress accelerates embryo development during seed development in oilseed rape plants remains to be elucidated. A deeper understanding of the effects of prolonged elevated temperatures on seed development could prove invaluable in the generation of thermotolerant canola varieties, which would help to improve crop yields in the context of global warming in the 21^st^ century.

## 2. MATERIALS AND METHODS

### 2.1 Plant materials and growth conditions

The *Brassica napus* spring cultivar Topas DH4079 was used in this study. The plants were treated as previously described (Mácová et al. 2022). Seeds were sterilised with 20% bleach, washed twice in sterile distilled water, vernalized at 4°C for 24 h, germinated on plates containing MS medium for 5 days (21°C, 16/8 h long-day photoperiod, LED light with light intensity of 150 μmol.m^-2^.s^-1^). Seedlings were transferred to soil in 0.7 L pots and grown for two weeks in the phytotron (21°C, 16/8 h long-day photoperiod, LED light with light intensity of 150 μmol.m^-2^.s^-1^) with one fertilization treatment with KRISTALON™ Start [N-P-K (19-6-20) + 3% Mg + 7.5%S]. One week later, the plants were transferred to 1.5 L pots. At the emergence of the first flowering buds, plants were transferred to two greenhouse chambers (120 m^3^ cultivation space) with control temperature (CT) and heat (HT) conditions maintained under a long-day regime (16/8 h), LED lights with a light intensity of 150 μmol.m^-2^.s^-1^, 35%–45% humidity and temperature of 18°C during the night. During the day, from 6 am daily, the temperature gradually ramped up to reach 21°C (CT) at 8 am and 32°C (HT) at 11 am and ramped down from 8 pm (CT) and 6 pm (HT) to reach 18°C at 10 pm. The plants spent 14 h at 21°C (CT) or 7 h at 32°C (HT) daily. The temperature was uniformly distributed in the two chambers by an air circulation system. Temperatures were controlled with a thermostat in the middle of each chamber, situated on the wall at 150 cm from the ground, and adjusted by water heaters and an adiabatic cooling system. Plants were well watered manually in the trays to avoid any effect associated with drought stress. Plants were once fertilized with KRISTALON™ Fruit and Flower [N-P-K (15-5-30) + 3% Mg + 5%S] at the flowering start and kept in the respective chamber till seed harvest. Seeds used for the experiments were obtained from hand-pollinated flowers. The age of the seeds was counted from the day after pollination (DAP). Phytotrons and Greenhouse chambers (Photon Systems Instruments s.r.o., Czech Republic) were maintained by the technical personal of the Plant Sciences core facility of CEITEC Masaryk university, Brno, Czech Republic.

### 2.2 Seed phenotyping

Roughly ten hand-pollinated and dried siliques from four plants in each growth condition per biological replicate, repeated five times, from CT (n = 116 siliques in total) and HT (n = 245 siliques in total) conditions were screened for the total, intact (round intact seeds with intact embryo), seeds undergoing SCR (partial or fully ruptured seed coat with intact embryo or germinating embryo respectively) and dead seeds (empty and shrunken without embryo) (Fig. S1a-c). Fresh seeds from 14 to 20 DAP were collected from Topas grown at CT and HT for embryo staging. The whole seed and its isolated embryo were imaged using a Levenhuk M500 base digital camera attached to a Zeiss stemi 305 stereomicroscope. Later, the images were used to measure the seed and embryo areas using ImageJ for 14, 18 and 20 DAP seeds. The ratio of the embryo-to-seed area was calculated to get information on the space occupied by the embryo within the seed at each stage of development. Statistical significance was accounted for by a two-way ANOVA test with Tukey’s multiple comparisons.

### 2.3 RNA extraction and sequencing

RNA was extracted from 50 mg of ground tissue from intact 14, 18 and 20 DAP seeds under CT and HT conditions, and from 20 DAP SCR seeds grown under HT conditions. Whole seeds were collected from three plants in batches of four replicates, per developmental stage, per growth condition. using a NucleoSpin RNA Plant and Fungi kit (Macherey-Nagel) following the manufacturer’s protocol. The samples were treated with RNase-free rDNase (Macherey-Nagel) and repurified using the RNeasy MinElute Cleanup Kit (Qiagen). The samples were sequenced by Novogene Genomic Sequencing Labs. After sample quality control, mRNA library was prepared after poly-A enrichment using NEBNext Ultra II Directional RNA Library Prep Kit for Illumina. Samples were sequenced using paired-end short-read (75 bp) on an Illumina NovaSeq Sequencing platform PE150 with sequencing depth ranging from ~62M reads in sample THT_14D_4 to ~92M reads in sample TCT_14D_2 (Table S1). Quality check of raw paired-end fastq reads was carried out by FastQC (Andrews 2010), and their origin was categorised using BioBloomTools v2.3.4 (Chu et al. 2014). The Illumina adapters and quality trimming of raw fastq reads were performed using Trimmomatic v0.39 (Bolger et al. 2014). After trimming, a mean of 71.1M ± 8.0M reads per sample were retained (range: 62.3M-92.2M), corresponding to more than 99% of the raw reads in all samples. Trimmed RNA-Seq reads were mapped to the *Brassica napus* genome (genome build: Bra_napus_v2.0) with gene annotation (NCBI *Brassica napus* Annotation Release 101) using STAR v2.7.3a (Dobin et al. 2013) as splice-aware short read aligner and default parameters except outFilterMismatchNoverLmax 0.1 and twopassMode Basic. Quality control after alignment concerning the number and percentage of uniquely-and multi-mapped reads (Table S1), rRNA contamination, mapped regions, read coverage distribution, strand specificity, gene biotypes, and PCR duplication was performed using RSeQC v4.0.0 (Wang et al. 2012)(Wang et al. 2012), Picard toolkit v2.25.6 (Broad Institute and GitHub Repository 2018), Qualimap v.2.2.2 (Okonechnikov et al. 2016). All compatible results and statistics were processed by MultiQC v1.10.1 (Ewels et al. 2016). The sequence data have been deposited in the NCBI under the Bio-Project PRJNA1188476 (https://www.ncbi.nlm.nih.gov/bioproject/PRJNA1188476).

The differential gene expression analysis was calculated based on the gene counts produced using featureCounts from Subread package v2.0 (Liao et al. 2014) and further analysed by Bioconductor package DESeq2 v1.34.0 (Love et al. 2014). DESeq2 with independent filtering was used to control false discovery rate and chosen for the differential gene expression analysis. Difference between compared samples was expressed in Log2FC calculated by comparing the average normalized counts of individual genes from one condition versus another condition, followed by log2 transformation. Genes were considered to be differentially expressed based on a cut-off for adjusted p-value ≤ 0.05 and |log2FC| ≥1, applied to shrunken fold changes obtained via lfcShrink() using the apeglm method (Zhu et al. 2019) The data are compiled in Table S1 (RNAseq sample description), Table S2 (Full dataset for the DEGs for all the comparisons), Table S3 (Differentially regulated GO terms associated with plant cell wall, cell cycle and developmental pathways), Fig. S8 (Number of shared downregulated DEGs), and Fig. S9 (Number of shared upregulated DEGs). Clustered heatmaps were generated from selected top differentially regulated genes using R package pheatmap v1.0.12 (Kolde 2015), volcano plots were produced using ggplot2 v3.3.5 package (Wickham 2011), and MA plots were generated using ggpubr v0.4.0 package (Kassambara 2018). GO enrichment analysis was performed using the Bioconductor package clusterProfiler v3.12.0 (function enricher) with the Gene Ontology database format version 1.2, and data version releases (2023-05-10). Gene IDs were converted to UniProt IDs through three complementary approaches: (1) direct cross-mapping of Entrez gene IDs against UniProt IDs; (2) BLAST alignment of transcript sequences against the NCBI Nucleotide database restricted to taxon 3701 (*Arabidopsis* genus), with resulting RefSeq IDs cross-mapped to UniProt; and (3) the same BLAST approach extended to include taxon 981071 (*Brassicaceae* tribe). The background gene universe was defined as the union of all successfully converted IDs across these three approaches (~48,000 genes from an initial set of ~84,000 per DESeq2 results table, representing a loss of ~33% during ID conversion). Multiple testing correction was applied using the Benjamini-Hochberg (BH) method. A q-value threshold of 1 was used to retain all enrichment results for subsequent manual filtering.

### 2.4. Seed embedding and sectioning for seed coat thickness measurements

Seeds were fixed in FAA (45% ethanol, 5% acetic acid and 1.85% formaldehyde) for 24 h under vacuum, followed by another incubation at 4°C for 24 h in fresh FAA. The seeds were washed once with water. Tissue softening was done with Franklin Maceration solution (1:1 v/v of 30% H_2_O_2_ and glacial acetic acid) for 4 h at 60°C. Dehydration was done using 10, 20, 30, 40, 50, 60% ethanol series in 10% glycerine for 1 h at 30°C, followed by 4 days in 70% ethanol at 4°C and dehydration with 80, 90 and 95% ethanol for 1 h at room temperature. Infiltration and embedding with Technovit 7100 (Kulzer Technik GmbH) were performed according to the manufacturer’s protocol with the following modifications. The initial infiltration was performed in a 50% infiltration solution in ethanol overnight at room temperature, followed by incubation for one week at 4°C in 100% infiltration solution with 1% w/v Technovit 7100 Hardener I. Embedding was performed with 6.6% w/v Technovit 7100 Hardener II in infiltration solution and incubated for 8 days at −18°C to delay initial polymerisation. The samples were then incubated at room temperature to complete the polymerisation process. The 3-µm sections were prepared with a Leica RM2235 microtome using an attached stainless-steel knife.

### 2.5 Toluidine blue staining

Ten seeds were pooled from three independently grown plants per condition per developmental stage. The seeds were used as operational unit of replication. The sections were stained with 0.05% Toluidine blue (Acros Organics) for 1 minute, followed by four washing steps of three minutes each with water. The slides were then dried on the hot plate maintained at 70°C and mounted in Entellan (Sigma-Aldrich). Imaging of the sections was performed using a Zeiss AxioImager microscope. Measurements of the thickness of seed coat layers from twenty regions of interest (ROI) were averaged into a single value per section. The values from three sections per seed were averaged into a single value. The thickness was measured using ImageJ. Statistical analysis (two-way ANOVA test with Tukey’s multiple comparisons) was performed at the level of individual seeds (n = 10 seeds per developmental stage per condition) to compare the thickness of the seed coat at the seed level.

### 2.6 Staining of lignin and pectin

Seven seeds per condition per developmental stage were pooled from three independently grown plants from a single growth round. For this experiment, 14 and 18 DAP from CT and HT and 22 DAP only from CT (for comparison with 18 DAP from HT) were used. Pectin was detected using a 0.05% w/v Ruthenium Red stain (Sigma-Aldrich) in an aqueous solution. Sections were rehydrated, stained for 7 minutes in Ruthenium Red, followed by two wash steps with water, drying on the hot plate and mounted in Entellan (Sigma-Aldrich). Lignin was detected with 0.5% Safranin-O (Acros Organics) in an aqueous solution; stained for 20 seconds followed by three washes in water. Slides were mounted in Entellan (Sigma-Aldrich) after drying. Imaging was done using a Zeiss AxioImager microscope.

### 2.7 Immunolabelling

Seeds at 10, 14, and 18 DAP from CT and HT, in two biological replicates, were fixed in 4% w/v paraformaldehyde in ice-cold PEM buffer (50 mM PIPES, 5 mM EGTA, and 5 mM MgSO_4_, pH 6.9). Samples were incubated on ice under vacuum for twice 1 h with fresh fixative solution, followed by overnight incubation at 4°C. The samples were washed twice with ice-cold PEM buffer, dehydrated through an ethanol series (50%, 70%, 80%, 90% and 100% three times) under vacuum and infiltrated with increasing concentrations (30%, 50% and 100%) of LR White resin (Agar Scientific) in absolute ethanol over 8 days before being polymerised at 60°C for 24 h. Six seeds per DAP per growth condition per biological replicate were sectioned (1 µm thickness) using a glass knife mounted on a Leica RM2265 or EM UC7 microtome. Immunolocalizations were performed with JIM5 (anti-Pectic polysaccharide, binding to partially methylesterified and unesterified homogalacturonan, monoclonal, AS18 4194, Agrisera), LM19 (anti-Pectic polysaccharide, binding to unesterified homogalacturonan, monoclonal, AS18 4191, Agrisera), LM20 (anti-Pectic polysaccharide, binding to methylesterified homogalacturonan, monoclonal antibody, Agrisera AS18 4193), and LM25 (anti-Xyloglucan, monoclonal, AS18 4205, Agrisera) antibodies. For immunolocalization with LM25, an incubation with 0.1% pectolyase in 1X PBS was done for 25 min at room temperature and washed once in 1X PBS before the blocking step. For immunolocalization with JIM5, LM19, and LM25, the sections were blocked with 3% w/v BSA in 1X PBS and incubated for 1 h at room temperature. For the immunolocalization with LM20, an incubation in CAPS buffer (50 mM CAPS and 2 mM CaCl_2_) for 1 h followed by three washes with 1X PBS was performed before blocking with 3% skimmed milk in 1X PBS. Sections were incubated overnight at 4°C in a humid chamber with antibody diluted 1:10 v/v in 1% BSA in 1X PBS for JIM5, LM19 and LM25, or in 1% skimmed milk in 1X PBS for LM20. The sections were brought to room temperature for 1 h, washed three times with the above-mentioned buffers and incubated for 1 h in a humid chamber in the dark with the secondary antibodies. A dilution of 1:100 v/v of secondary antibodies [anti-rat IgG Alexa 488 (Thermo Fisher) or anti-rat IgG Dylight 488 (Agrisera) for JIM5 and anti-rat IgM Dylight 488 (Thermo Fisher) for LM19, LM20 and LM25] in 1% BSA or 1% skimmed milk in 1X PBS was used. Sections were washed in the respective buffers and 1X PBS, counterstained with 0.025 mg/ml Calcofluor White (Sigma-Aldrich) and mounted with VECTASHIELD antifade mounting medium (Eurobio). The stacked images were obtained with the ZEISS Axioimager 2 with a 40X dry objective.

The intensity of the immunofluorescence in the cell walls of the epidermal and palisade layers of the seed coat was quantified using ImageJ software on unadjusted raw images. For the three epidermal layers, a segmented line with a width of 25 points was used to manually mark the cell walls of adjacent cells of the same layer (selection of ROIs using the Calcofluor White image). The sum of fluorescence intensity from the immunolocalization signal on the ROIs was represented as one measurement for each of the three epidermal layers. A ROI was marked using the polygon selection tool for the palisade layer. The fluorescence value obtained over the ROIs for the different layers was subtracted from the background signal over the same area. Finally, normalised fluorescence intensity was calculated and used for statistical testing. Images were adjusted using min/max settings for optimal representation of fluorescent signals in the figures.

### 2.8 Cell wall profiling from the seed coat

The enzymatic fingerprinting of the cell wall of the seed coat from 10, 14 and 18 DAP seeds grown under CT and HT conditions was performed in three biological replicates (independent batches of plants). For each replicate, the seed coats were dissected from the 20 seeds per developmental stage per condition and were submerged in 96% (v/v) ethanol. The ethanol fraction was discarded, and the seed coats were dried in a speed vacuum concentrator at 30°C overnight. Dried seed coats were digested with 5 U/mg sample dry weight of a mixed solution of *Pectobacterium carotovorum* endo-polygalacturonase and *Paenibacillus* sp. xyloglucanase (Megazyme, Bray, Ireland) in 50 mM ammonium acetate buffer (pH 5) at 37°C for 18 h. Samples were centrifuged at 13,000 rpm for 10 min, and 100 µl of the supernatants were transferred into vials. For MS analysis, 10 μl of each fraction was injected. The oligosaccharides released from digestion were separated according to Voxeur et al. (2019). Blank samples consisted of enzyme solution and buffer, while quality control samples consisted of citrus pectins with enzymes and tamarind xyloglucans with enzymes. Xyloglucan and pectin fragments were analysed using an HPLC system (UltiMate 3000 RS HPLC system, Thermo Scientific, Waltham, MA, USA) coupled to an Impact II Ultra-High Resolution Qq-Time-Of-Flight (UHR-QqTOF) spectrometer (Bruker Daltonics, Bremen, Germany) equipped with an electrospray ionisation (ESI) source in negative mode with the end plate offset set voltage to 500 V, capillary voltage to 4000 V, nebuliser to 40 psi, dry gas to 8 l/min and dry temperature of 180°C. The Compass 1.8 software (Bruker Daltonics) was used to acquire the data. Chromatographic separation was performed on an ACQUITY UPLC Protein BEH SEC Column (125A, 1.7 μm, 4.6 mm x 300 mm, Waters Corporation, Milford, MA, USA) coupled with a guard column BEH SEC Column (125A, 1.7 μm, 4.6 mm x 30 mm). Elution was performed in 50 mM ammonium formate and 0.1% formic acid at a flow rate of 0.4 ml/min and with a column oven temperature of 40°C. Mass accuracy was used for feature annotation with a threshold of 0.005 Da, retention time was matched within 0.2 min and MS/MS confirmation was not used for peak identification.

### 2.9 Space-restricted growth of siliques

Silicone tubes (length 5.0 cm, outer diameter 5.0 mm, wall thickness 1.0 mm, inner diameter 3.0 mm, hardness 60 Shore A; P-lab, cat. no. L905756) were inserted over developing siliques at 10 DAP and 12 DAP under HT and CT conditions, respectively, in four biological replicates of three plants for each condition using ~ 20 siliques per replicate. A 10 DAP silique at HT has a similar silique thickness to a 12 DAP silique at CT. Siliques from the same growth conditions without a silicone tube were used as negative control. The number of total, intact, dead and SCR seeds per silique were estimated when mature and fully dry. For seed coat thickness measurements of space-restricted seeds, seed embedding with LR White and seed sectioning (protocol described in section 2.7) was used followed by toluidine blue staining (protocol described in section 2.5). For seed coat thickness measurements, and seed and embryo size measurements, seeds at 24 DAP from CT and 20 DAP from HT were used because they were at the same developmental stage.

### 2.10 Nanoindentation

A cylindrical 3D-printed mould of 2-cm diameter and 0.5-cm height was created with multiple circular wells of 1.8 to 2.4 mm diameter and 1.5-mm depth on the surface for holding the Topas seeds in place during the measurements (Fig. S7). Three fresh 24 DAP seeds from CT and three fresh 20 DAP seeds from HT conditions were used for this study. The seeds were kept hydrated in a vial contained a wet tissue paper. A Hysitron TI 950 instrument (upgraded to TI980) equipped with a diamond conical indenter with a cone angle of 90° and a tip radius of 0.97 µm was used for the nanoindentation. The indenter was calibrated on fused silica standard and tested on a polycarbonate standard sample. Accelerated mechanical property mapping (XPM) was applied, as this mode enables the acquisition of elastic modulus maps with high spatial resolution and short acquisition times. Compared to the classical quasistatic indentation method, XPM is substantially faster, the time required to perform a single quasistatic indent is more than one order of magnitude longer. The reduced measurement time minimizes the effects of thermal drift, seed drying and suppresses the time-dependent mechanical response of the seed material. The maximum load was set to 50 µN, with a loading time of 0.5 s, a holding segment at the maximum load of 0.5 s, and an unloading time of 0.1 s. Each measurement consisted of a 7 × 7 array of indents covering an area of 7 × 7 µm^2^. In total, more than 150 nanoindentation data points were acquired using this method. The data were evaluated according to Oliver and Pharr (1992). The indentation reduced modulus values were acquired directly from the analysis of the indentation unloading curves (Fig. S10). The reduced modulus *E*_r_ is calculated as a combination of the sample material and indenter elastic deformations:

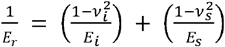

Here, *E*_*s*_, *V*_*s*_ and *E*_*i*_, *V*_*i*_ are the Young’s moduli and Poisson ratio values of samples and the diamond indenter, respectively. If the studied samples exhibit low elastic moduli compared to diamond, the first term in the equation may be neglected.

The hardness *H*_*IT*_ was calculated from the ratio of applied load and the contact area of the indenter and the sample.

### 2.11 Statistical analysis

The number of seeds used for analysis in each experiment is chosen accounting the time and material constraints. GraphPad Prism v.8.4.3 was used to plot the graphs and calculate the statistical significance. For the RNA sequencing experiment, R programming was used to analyse, represent data and obtain the statistical significance. Asterisks indicate significant difference (*, **, *** and **** correspond to p-values of 0.05 > p > 0.005, 0.005 > p > 0.0005, 0.0005 > p > 0.00005 and p < 0.00005) unless otherwise stated. Seed phenotype counts were obtained from siliques collected from four plants per condition across five biological replicates (total number of scored siliques: CT n = 116, HT n = 245) and are presented both as percentages and as actual counts per silique. Statistical significance was assessed using a quasi-binomial generalized linear model (logit link) fitted to the seed counts per silique, with each phenotypic category (intact, SCR, dead) modelled as the number of seeds in that category out of the total per silique; the quasi-binomial family was used to account for overdispersion among siliques. Embryo and seed area measurements were analysed by a two-way ANOVA test (temperature × developmental stage) with Tukey’s multiple comparisons test for assessing statistical significance (n = 30 per condition per stage). Seed coat layer thickness was analysed by two-way ANOVA (temperature × developmental stage) followed by Tukey’s multiple comparisons test, with statistical testing performed at the level of individual seeds (n = 10 per condition per stage). Results of the pairwise comparisons are represented as a letter, in which groups sharing a common letter do not significantly differ (p ≥ 0.05). Immunofluorescence intensity was quantified from three sections averaged per seed, with six seeds per condition per developmental stage from each of two biological replicates. Statistical significance was assessed by two-way ANOVA (temperature × developmental stage) followed by Tukey’s multiple comparisons test, performed at the level of individual seeds (n = 12 per condition per stage across both biological replicates). Enzymatic fingerprinting of the seed coat cell wall was performed in three biological replicates per condition per developmental stage, and oligosaccharide abundance is expressed as MS peak area. Statistical significance was assessed by two-way ANOVA (temperature × developmental stage) followed by Tukey’s multiple comparisons test (n = 3 biological replicates). Nanoindentation was performed on three seeds per condition (20 DAP HT and 24 DAP CT, matched for developmental stage), with multiple indents acquired per seed and averaged to a single reduced modulus (E_r_) and hardness (H_IT_) value per seed. Statistical significance between conditions was assessed at the level of individual seeds (n = 3) using Student’s t-test.

## 3. RESULTS

### 3.1 *Brassica napus* seed coat integrity is affected by long-term heat stress

We have previously reported that in *Brassica napus* cv. Topas early seed development is affected by long-term heat treatment (HT) (Mácová et al. 2022). This study focuses on the seed coat rupture (SCR) observed in seeds of Topas developed under HT (Fig. 1b), starting at 20 DAP, while this phenomenon has very rarely been observed in seeds grown under control treatment (CT) (Fig. 1a, Fig. S1). The SCR phenotype occurred with variable outcomes, ranging from a minor or incomplete break of the seed coat with intact tegmen and embryo remaining within the seed to a complete seed coat rupture resulting in the embryo protrusion through the break site appearing like a germinated seed (Fig. 1b, Fig. S1d-g). Such embryos often had distinguishable patterning defects induced by high temperatures: fused cotyledons, cotyledons with asymmetric shape, only one or three cotyledons, aberrant root tip (Fig. S1g). The quantification of the seed phenotypes by counting the number of dry seeds per silique revealed a significant decrease of intact seeds, accompanied by a significant increase in the proportion of dead seeds and SCR seeds for plants developed under HT compared to that of CT (Fig. S1h). At HT, approximately 50% of the seeds in a silique underwent SCR (Fig. 1c).

### 3.2 The embryo-to-seed size ratio is increased at HT

Embryo staging of the seeds from 14 to 20 DAP revealed that embryo development is accelerated in seeds developed under HT compared to seeds grown under CT (Fig. 1d), which is consistent with our previous observations in seeds at early developmental stages (Mácová et al. 2022). We visually estimated that the embryos in HT-grown seeds were approximately four days ahead of embryos in CT-grown seeds in their developmental stage, e.g., a 14 DAP embryo at HT would be developmentally similar to an 18 DAP embryo at CT (Fig. 1d).

We studied the seed area and embryo area of seeds from 14 to 20 DAP at CT and HT. Embryos at HT were more than twice the size of those at the same DAP at CT (Fig. 2a). The seed area increased from 14 to 20 DAP, and this increase slowed by 20 DAP as seeds entered maturation (Fig. 2b). The HT seeds grew faster than CT seeds between 14 and 20 DAP. Indeed, the seed size at 18 DAP was significantly larger under HT than under CT (Fig. 2b), whereas there was no significant size difference at 14 DAP and 20 DAP. We noticed that the seeds did not undergo a compensatory increase in size to accommodate the HT-induced larger embryos. The embryo-to-seed area ratio provides valuable insight into the space occupied by the embryo within the seed at a given DAP under the two growth conditions (Fig. 2c). We observed that the embryos developed under HT filled significantly more space in the seed than those grown under CT. These observations suggest that the accelerated embryo growth in HT resulted in the formation of larger embryos but not larger seeds.

**Fig. 2.**
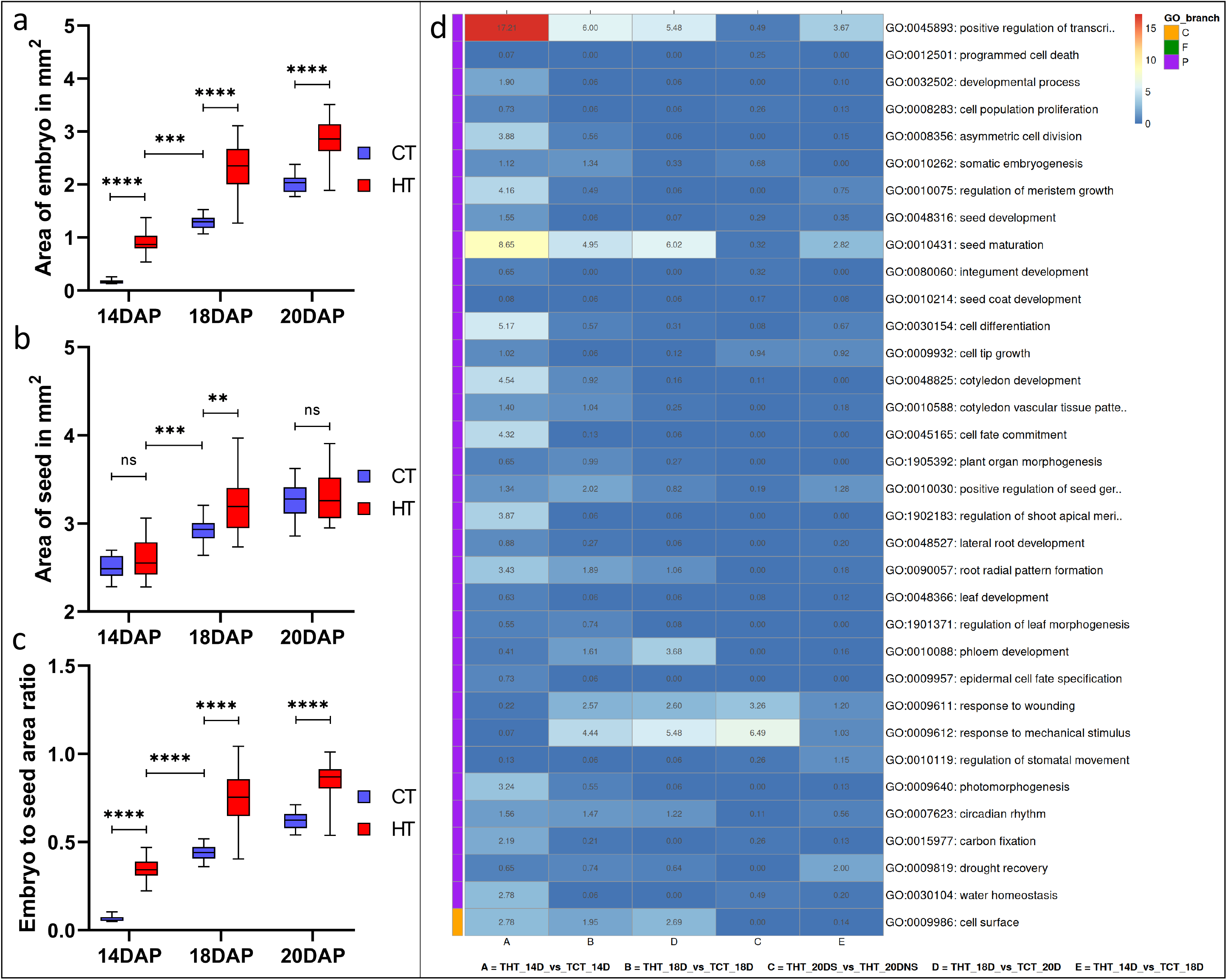
Embryo and seed size measurements of *Brassica napus* cv. Topas seeds developed under CT and HT. (a, b) Embryo (a) and seed (b) area measurements in mm^2^ of 14, 18 and 20 DAP Topas seeds developed under CT (blue) and HT (red). (c) Ratio of embryo-to-seed size in 14, 18 and 20 DAP seeds developed under CT and HT. Asterisks indicate statistically significant differences in HT and between 14 DAP HT and 18 DAP CT (*, **, *** and **** correspond to *p*-values of 0.05 > *p* > 0.005, 0.005 > *p* > 0.0005, 0.0005 > *p* > 0.00005 and *p* < 0.00005, respectively); n.s., not significant. (d) Upregulated GO terms (Log_2_FC≥1) related to plant development with the transcriptional response of Topas seeds to HT (values in the box depict –Log_10_(q-value)).

When seed and embryo size were compared at the same embryo developmental stage, 14 DAP HT seeds and embryos were smaller than 18 DAP CT seeds.

As the embryonic developmental shift could be observed phenotypically, we investigated the transcript profiles by RNA-seq of these seeds (14, 18, 20 DAP) developed in CT and HT. A Principal Component Analysis (PCA) (Fig. S2) indicated that the 14 DAP HT seeds clustered together with the 18 and 20 DAP CT seed samples, further validating the observations of the HT-induced accelerated seed developmental process. Next, we screened the gene ontology (GO) enrichment analysis for GO terms associated with the process of seed development and cell cycle regulation (Fig. 2d, Fig. S3, Table S3). A significant enrichment of GO terms associated with seed maturation and positive regulation of transcription was found when comparing the HT and CT datasets from seeds of the same development stage (14 DAP HT vs. 18 DAP CT). GO terms associated with DNA biosynthesis, replication and mitotic DNA replication initiation are significantly enriched comparing the datasets from SCR and non-SCR 20 DAP seeds developed at HT (Fig. S3, Table S3).

### 3.3 The formation of larger embryos at HT impacts the thickness of the seed coat cells

We hypothesised that accelerated embryo growth, together with a potential premature transition into the seed maturation phase, may disrupt the normal development of the maternally-derived seed coat, thereby contributing to the SCR phenotype. Notably, the seed coat (maternal origin) and the embryo (zygotic origin) differ both genetically and in their developmental timing, influenced by high temperature. These fundamental differences should be considered when interpreting whole-seed RNA-seq data, as signals from distinct tissues with asynchronous developmental programs may mask tissue-specific transcriptional dynamics. To further explore these effects, we next examined how HT impact seed coat morphology. The seed coat of *B. napus* is composed of three layers of cells (Fig. 3a): an outer epidermal layer of three cell files, a middle palisade layer made of a single layer of cells with substantial secondary cell walls, and an inner endothelial layer surrounding the endosperm (Jiang and Deyholos 2010). The thickness of these three layers of the seed coat was measured in seeds of 10, 14, 18, 20, and 22 DAP developed under CT and HT (Fig. 3). All three seed coat layers were significantly thinner in 14 to 22 DAP seeds developed under HT (Fig. 3b-h). Taking into account the same embryonic developmental stage, we compared the seed coat thickness of seeds from 18 DAP at CT with 14 DAP at HT and 22 DAP at CT with 18 DAP at HT and observed a significant reduction in thickness of the three layers (Fig. 3b-e). The palisade and the first endothelial layers were the most affected, with a 46% reduction in the thickness of endothelial cell layer induced by HT as the seed aged from 14 DAP to 20 DAP, compared to only 12% at CT (Fig. S4a). These results indicate that the HT-induced accelerated embryo growth between 14 and 20 DAP may physically affect the development of maternally-derived seed coat cells, leading to the thinning of seed coat layers. We hypothesised that faster embryonic growth might increase the strain in the seed coat cells. In this context, we performed a detailed examination of cell wall constituents in the seed coat.

**Fig. 3.**
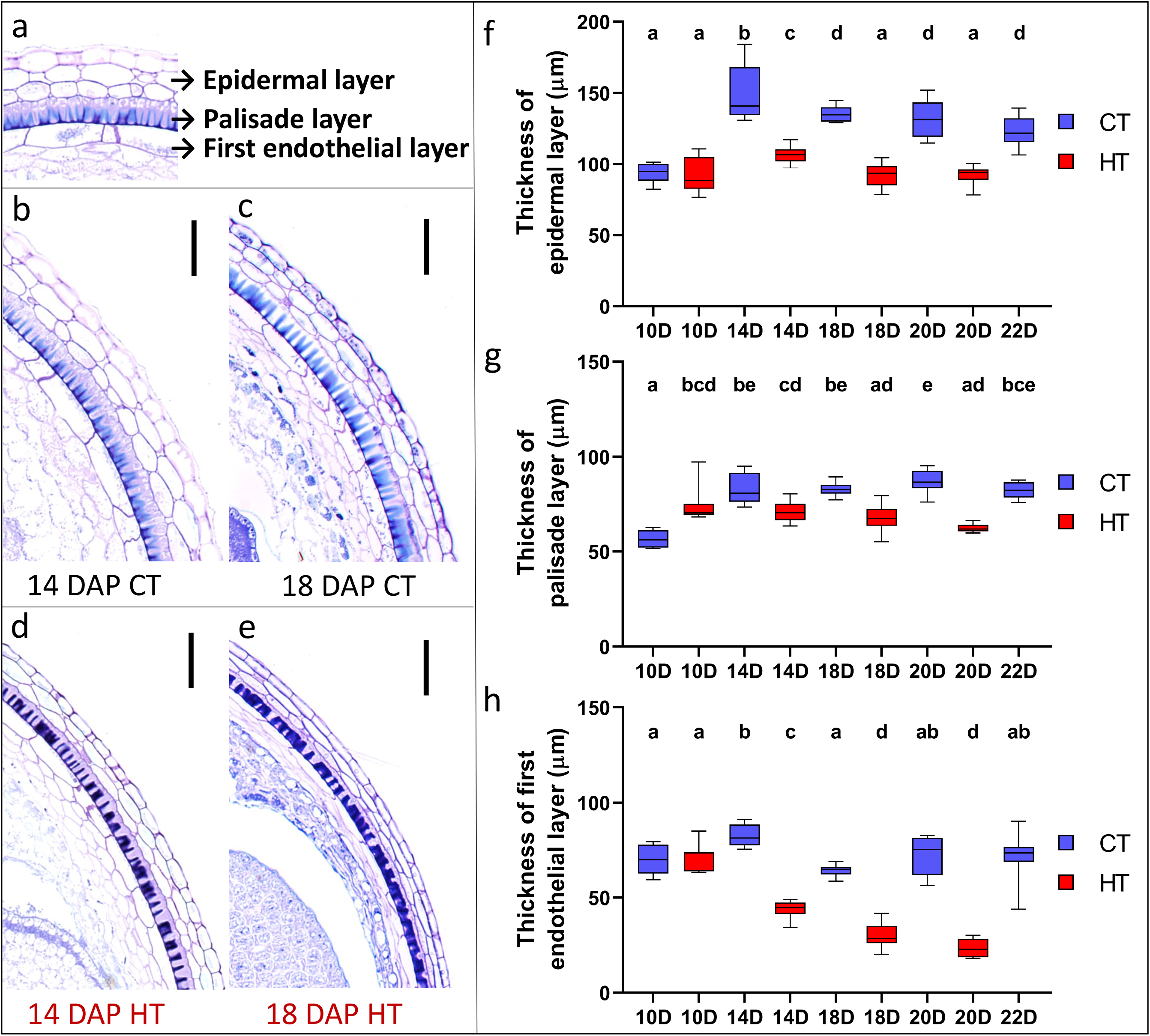
The thickness of the seed coat cells is reduced under HT. (a-e) Histological sections of Topas seeds focused on the seed coat, stained with toluidine blue. (a) The seed coat is composed of an epidermal layer, a palisade layer and an endothelial layer. (b-e) Seed coat of 14 DAP seeds from CT (b) and HT (d) and 18 DAP seeds from CT (c) and HT (e). Scale bars represent 200 µm. (f-h) Box-whisker plots show the distribution of seed coat layer thickness in µm and the box indicates the interquartile range, the central line for median and the Tukey whiskers of the epidermal layer (f), palisade layer (g) and first endothelial cell layer (h) in 10, 14, 18, 20 DAP, and 22 DAP CT seeds (blue) and 10, 14, 18, and 20 DAP HT seeds (red) (n = 1-seeds per DAP per condition). Samples with the same letter do not statistically differ.

### 3.4 The composition and properties of the seed coat cell wall are altered in HT-grown seeds

Given the developmental differences between CT and HT conditions, we compared whole-seed RNA-seq datasets based on embryo morphology rather than seed age, aligning 18 DAP control (TCT_18D) with 14 DAP HT (THT_14D) samples. While this approach enables comparison of seeds with embryos at similar developmental stages, it inherently introduces confounding factors. Consequently, the observed transcriptional differences are unlikely to reflect developmental stage alone, but instead represent a combination of developmental, temporal, and temperature-induced effects. We also compared the transcriptome profiles of seeds exhibiting SCR (THT_20DS) and non-SCR (THT_20DNS) phenotypes. GO enrichment analysis revealed significant differences in the regulation of many pathways related to plant cell walls (Fig. 4a). These included GO terms related to cell wall biogenesis, cell wall thickening, and cell wall modification. Interestingly, the datasets of HT-grown seeds also showed over-representation of cellulose, pectin, and xyloglucan metabolism and modification pathways. Examining the differentially expressed genes (DEGs) associated with pectin metabolic pathways revealed differential regulation of enzymes such as galacturonosyltransferases (GAUTs), which are involved in pectin biosynthesis (Mohnen 2008), pectinesterases and pectinesterase inhibitors, involved in the chemical modification of pectin (Wormit and Usadel 2018; Wu et al. 2018; Xu et al. 2022), and polygalacturonases and pectate lyases, which are involved in pectin degradation (Marin-Rodriguez 2002; Park et al. 2010; Safran et al. 2023) (Fig. 4b).

**Fig. 4.**
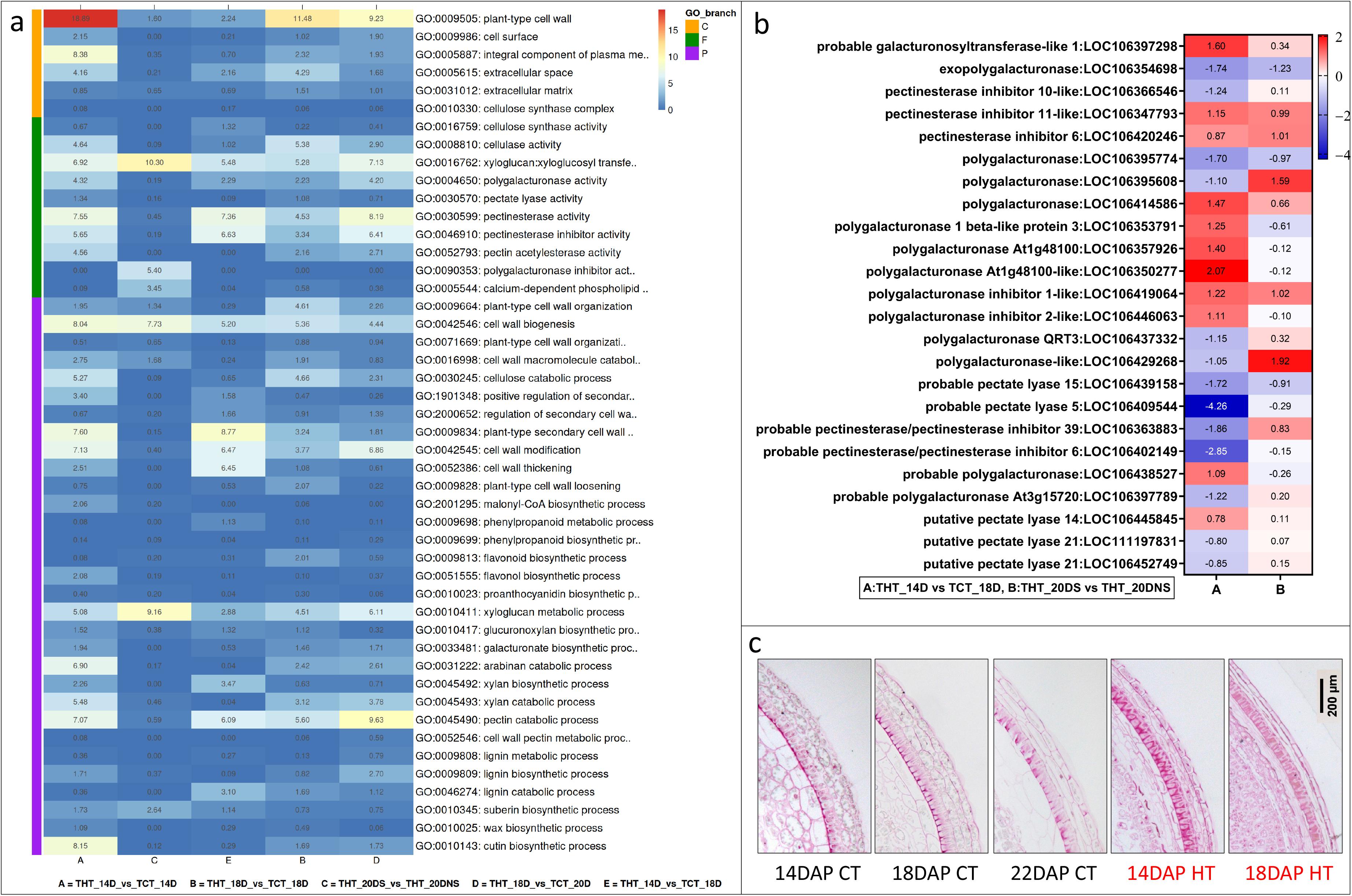
Transcriptional changes in Topas seeds associated with growth at HT indicate changes in cell wall composition. (a) GO terms associated with cell wall metabolism and regulation with the transcriptional response of Topas seeds to HT are shown (values in the box depict – Log_10_(q-value)). (b) Heatmap showing the differential transcription of selected genes involved in pectin metabolism in 14 DAP HT vs. 18 DAP CT seeds (A), and SCR vs. non-SCR seeds (B) at 20 DAP from HT. The full dataset is presented in Tables S2 and S3. (c) Histological sections of Topas seeds focused on the seed coat, stained with Ruthenium Red for pectin. Sections of 14, 18, and 22 DAP CT seeds and 14 and 18 DAP HT seeds are shown. Scale bars represent 200 µm.

The transcriptomic analysis suggested heat-induced alteration of cell wall structural dynamics within the seed. Based on our observations of reduced seed coat thickness and enlarged embryos under HT conditions, we hypothesised that the seed coat may be the main tissue affected by these transcriptional changes. Therefore, to validate and spatially resolve these findings, we employed a histological approach and assessed the levels of pectin and lignin in the seed coat of HT-grown seeds using Ruthenium Red (Fig. 4c), which stains acidic polysaccharides and pectin-rich regions red (Nagayama et al. 2019), and Safranin-O (Fig. S4b), which stains phenolic lignin compounds in bright red (Bond et al. 2008). The cells of the palisade layer in the seed coat of 14 and 18 DAP Topas seeds grown under HT appeared to stain more intensively red with both dyes than those of 14, 18 and 22 DAP seeds grown under CT, based on representative repeated observations. These observations suggest an altered accumulation of cell wall components in the seed coat under HT conditions.

### 3.5 HT is associated with increased pectin demethylesterification in the seed coat cell wall

We further investigated the presence of pectin and hemicellulose in the layers of the seed coat by immunostaining (Fig. 5, Fig. S5a). To detect pectin, which is primarily composed of methylesterified and demethylesterified homogalacturonan (HG), we used the LM19 antibody, which binds preferentially to demethylesterified HG, the LM20 antibody that binds preferentially to methylesterified HG, and the JIM5 antibody, which exhibits a high affinity for partially demethylesterified HG (Fig. 5) (Verhertbruggen et al. 2009). For hemicellulose, we focused on xyloglucans using the LM25 antibody (Fig. S5a). The seed coat of 14 and 18 DAP HT-developed seeds had significantly higher fluorescence levels of the LM19 and JIM5 antibody signals in the palisade layer, indicating an increase in demethylesterified HG in HT seeds (Fig. 5a,b). In contrast, no significant differences in fluorescence intensity for the LM20 antibody signal were observed in the palisade layer in the seed coat of CT and HT seeds (Fig. 5c). In the epidermal layer of HT-developed seeds, the JIM5 and LM19 antibody signal was significantly higher in 14 DAP seeds, while this increase occurred at 18 DAP in CT seeds (Fig. 5a,b). The LM20 antibody signal was significantly lower in 18 DAP HT seeds than in all other samples, indicating a reduction in the amount of methylesterified HG (Fig. 5c). These results suggest an early enrichment of pectin and a shift towards demethylesterified pectin in the cell walls of the palisade and epidermal cells in 14 DAP HT-grown seeds. LM25 fluorescence was significantly higher only in the palisade cells of 18 DAP HT-grown seeds compared to CT seeds, suggesting a slight effect of temperature on the amount of xyloglucan in the seed coat of these seeds (Fig. S5a).

**Fig. 5.**
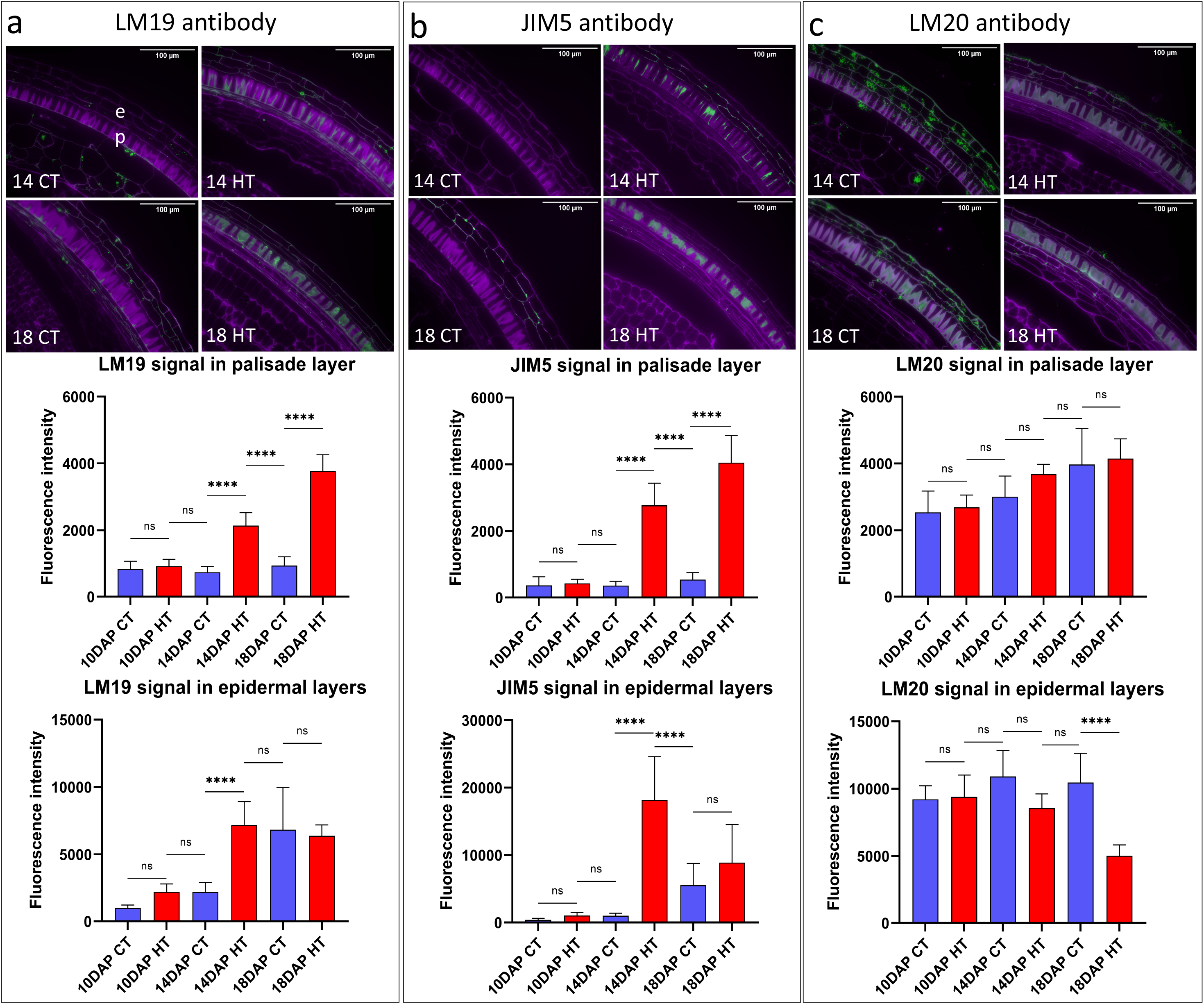
Immunolocalization to estimate the degree of pectin methylation using LM19 (a), JIM5 (b), and LM20 (c) antibodies on the seed coat of 14 and 18 DAP CT and HT seeds. The signal is visible in green; the cell structure is marked in magenta with Calcofluor White. Scale bars represent 100 µm. Fluorescence intensity was measured in epidermal (e) and palisade (p) cells in 10, 14 and 18 DAP HT and CT seeds. Asterisks indicate statistically significant differences between CT and HT and among seed ages (*, **, *** and **** correspond to *p*-values of 0.05 > *p* > 0.01, 0.01 > *p* > 0.001, 0.001 > *p* > 0.0001 and *p* < 0.0001, respectively); n.s., not significant.

To confirm our microscopic observations, we performed enzymatic fingerprinting of the cell wall of the isolated seed coats from 10, 14 and 18 DAP seeds of Topas grown at CT and HT (Fig. 6a). Endo-polygalacturonase hydrolyses HG preferentially in stretches of non-methylesterified galacturonic acid (GalA) residues, releasing several oligogalacturonans (OGs). OGs have been termed GalAxMeyAcz based on the degree of polymerisation of GalA and the number of methyl and acetyl groups. In our analysis of the Topas seed coat, we identified GalA3, GalA3ox, GalA3Xyl, GalA3Me, GalA3MeAc, GalA4Ac, GalA4Me, GalA4Meox, GalA4Me-H_2_O, and DPx>4Mey (methylesterified OGs with degree of polymerisation greater than 4). The total peak area obtained from the MS analysis was plotted by grouping OGs with and without methyl groups (Fig. 6b). A significant greater amount of demethylesterified OGs was released from the cell walls of 14 DAP seed coats at HT compared to CT, which correlates with the signal intensity of the LM19 antibody binding to the demethylesterified HGs in the cell walls in the seed coat layers (Fig. 6b,c). The release of oxidised OGs is also notable, as this may reflect the presence of long oxidised OGs trapped within the pectin and not solubilised through alcohol extraction. Regarding xyloglucan, LXXGAc levels remained unchanged and correlated with the LM25 signal in the seed coat (Fig. S5b,c). However, a significant increase in the level of fucosylated subunits was observed in the 14 DAP seed coats of HT compared to CT (Fig. S5b).

**Fig. 6.**
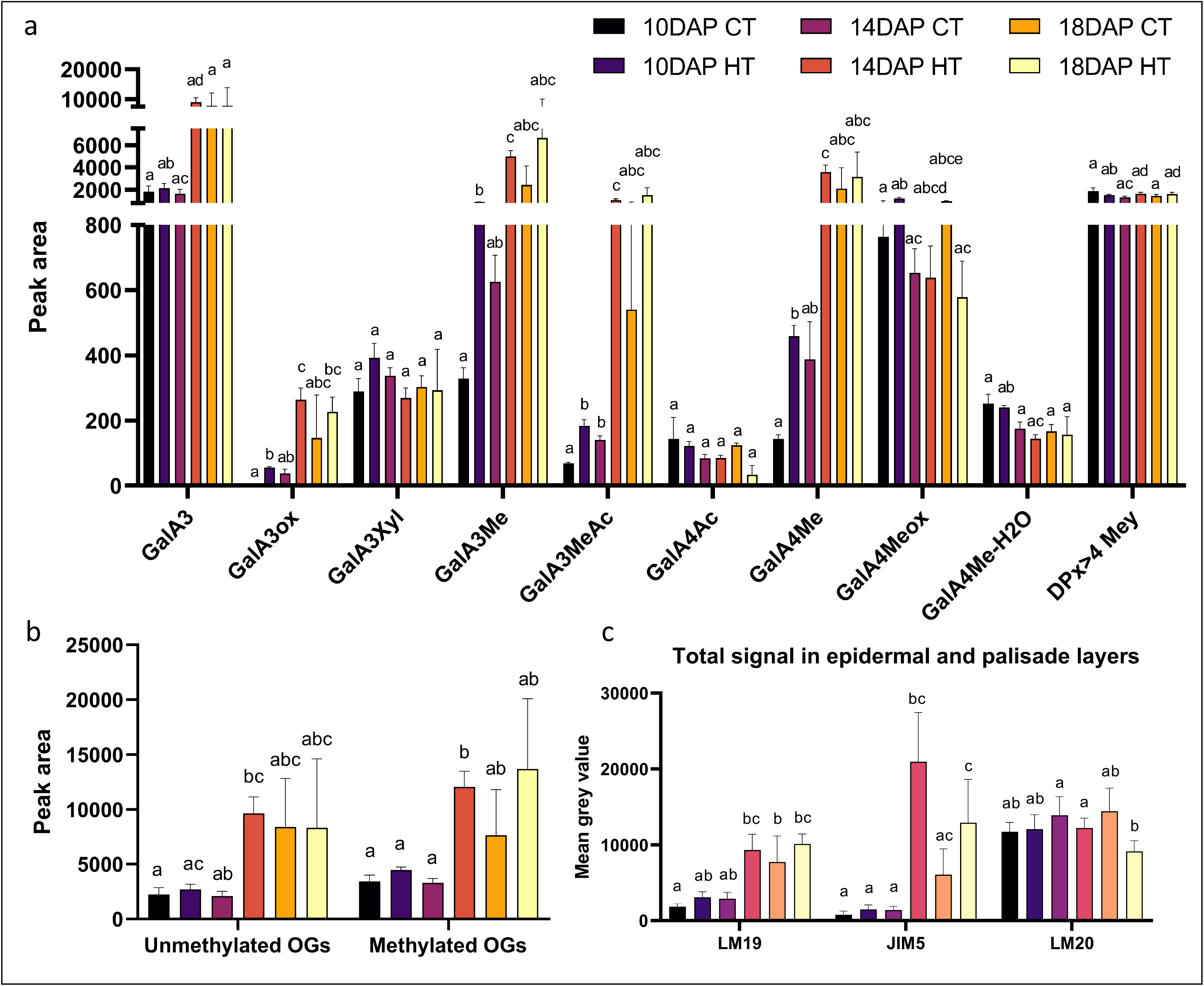
Pectin enzymatic profiling. (a) Pectin composition of the seed coat cell wall of 10, 14, and 18 DAP seeds from HT and CT. The graph shows the measurements expressed as peak area from the MS analysis for GalA3, GalA3ox, GalA3Xyl, GalA3Me, GalA3MeAc, GalA4Ac, GalA4Me, GalA4Meox, GalA4Me-H_2_O and DPx>4Mey (methylesterified OGs with degree of polymerisation greater than 4). (b) Graph showing the amount grouped as methylesterified versus non-methylesterified OGs from the data presented in panel (a). (c) Graph showing the total intensity of the fluorescence signal for pectin in the epidermal and palisade seed coat layers after immunolocalization using LM19, JIM5 and LM20 antibodies from Figure 5. The legend indicating the correspondence between colour and sample is the same for all three panels. Samples with the same letter do not significantly differ.

Using a variety of approaches, we uncovered that, between 14 and 18 DAP, seeds grew more at HT than at CT, probably due to the pressure exerted by embryo growth. We also found that growth at HT altered the properties of the cell walls in the seed coat of seeds from 14 DAP onwards. The cell walls of the epidermal and palisade cells accumulated more pectin and shifted prematurely towards producing demethylesterified pectin. These changes may have limited cell growth. Consequently, seeds grown at HT stopped growing after 18 DAP.

### 3.6 HT affects the physical properties of the seed coat

The composition of the seed coat cell wall determines its mechanical strength (Mendu et al. 2011; Sechet et al. 2018). The enrichment of GO terms associated with cell wall metabolism and the high content of demethylesterified pectin in the seed coat prompted us to investigate whether growth at a high temperature affects the mechanical strength of *B. napus* seed coat cell walls. We hypothesised that increased accumulation of demethylesterified pectin may alter cell wall mechanical properties, including stiffness and extensibility, and could potentially increase its susceptibility to rupture, thereby contributing to SCR. We analysed the mechanical properties of the seed coat cell wall using a nanoindentation technique, which provided valuable insights into the reduced modulus *E*_*r*_ (MPa) and the hardness *H*_*IT*_ (MPa). Reduced modulus quantifies the elastic response of the seed surface, e.g., its ability to return to its original shape after the release of the mechanical stress (Lapointe et al., 2025). It is a measure of stiffness, meaning the resistance of the seed to elastic deformation. The hardness measures the resistance to plastic (permanent) deformation and indicates how the seed surface may be resistant to permanent damage imposed by the mechanical stress. HT seeds exhibited a significantly lower reduced modulus *E*_*r*_ (Fig. 7a) and lower hardness *H*_*IT*_ (Fig. 7b) than CT seeds at the equivalent developmental stage (20 DAP HT vs. 24 DAP CT). Both the reduced modulus (*Er*) and hardness (*H*_*IT*_) indicated that the HT seed surface was mechanically more compliant and softer than that of CT seeds. Together with the reduced thickness of the seed coat layers (Fig. 3), these results indicate that the HT seed coat, as a whole structure, is mechanically weaker and less able to resist deformation than the CT seed coat.

**Fig. 7.**
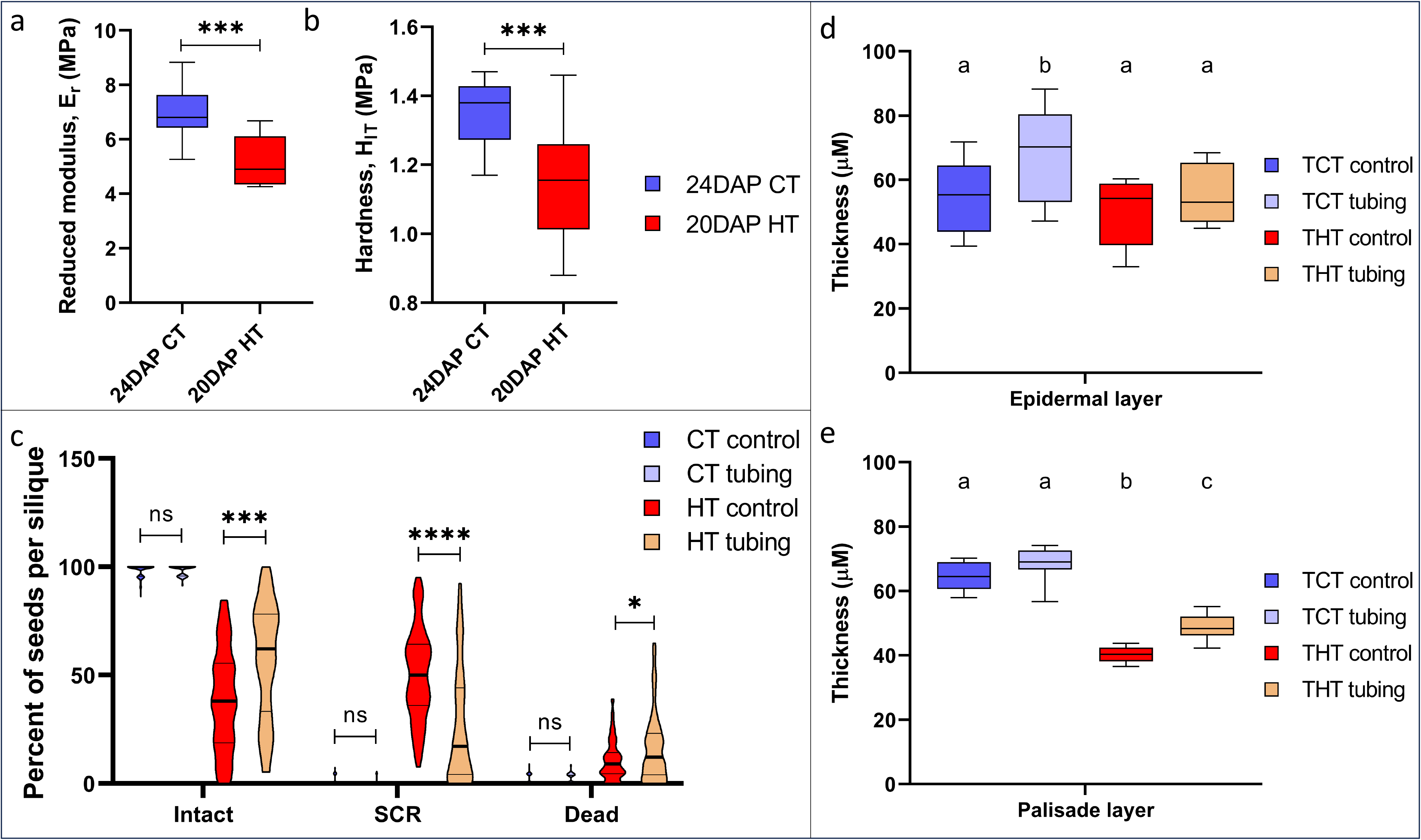
Seed coat mechanobiology of *B. napus* seeds. (a, b) Measurement of reduced modulus *E*_r_ (MPa) (a) and hardness, *H*_IT_ (MPa) (b), comparing 20 HT seeds and 24 DAP CT seeds. (c-e) A 3-mm silicone tube was applied to siliques from 12 DAP CT and 10 DAP HT. (c) Dry seeds were quantified for the percent of intact, SCR and dead seeds per silique. (d,e) Box-whisker plots show the size, median, outliers, and SD in µm of the thickness of the epidermal layer (c), palisade layer (d) of 24 DAP CT seeds and 20 DAP HT seeds to account for same developmental stage. The thickness of the endothelial layer was not measured in the tubing experiment. (a-c) Asterisks indicate statistically significant differences in HT and between seed ages (*, **, *** and **** correspond to *p*-values of 0.05 > *p* > 0.01, 0.01 > *p* > 0.001, and *p* < 0.001, respectively); n.s., not significant. (d,e) Samples with the same letter do not statistically differ.

### 3.7 Application of mechanical constraints during silique development partially rescues SCR phenotype

We found that the seed coat of HT-developed seeds had both a lower reduced modulus and lower hardness (Fig. 7a,b), consistent with a mechanically weaker and more compliant seed coat. We hypothesised that such a seed coat would be less able to withstand the pressure imposed by the faster-developing embryo, contributing to the occurrence of SCR. Should this be the case, then providing external support to oppose continuous embryo growth would prevent SCR. To test this hypothesis, we encased the developing siliques in a silicone tube to create an external mechanical constraint for the seeds, in an attempt to counteract the pressure exerted by the developing embryo on the seed coat (Fig. S6a). It has been demonstrated that applying a mechanical force to the siliques triggers seed maturation by arresting embryonic growth in *B. napus* Reston cv. (Rolletschek et al. 2024). In CT and HT seeds encased in silicone tubes, embryo size was significantly reduced, indicating that space restriction constrained embryo growth, whereas the ratio of embryo-to-seed area remained the same (Fig. S6c-e). It should be noted that the presence of a silicone tube may alter silique gas exchange, humidity, transpiration, local temperature, and maturation signalling. These factors were partially taken into account by using encased CT siliques as controls. The presence of the silicone tube on the siliques of CT plants did not affect seed viability or promote SCR (Fig. 7b, S6b). Under HT conditions, the external support provided by the silicone tube on the siliques resulted in a significant decrease in the number of seeds undergoing SCR and an increase in the number of intact seeds (Fig. 7c, S6b). Measuring the thickness of the seed coat layers of HT-grown seeds revealed that the palisade layer thickness increased significantly in seeds encased by the silicone tube compared to seeds developed in un-encased siliques (Fig. 7d,e). This external mechanical support may have partially compensated the reduced mechanical strength of the seed coat associated with faster embryo development.

## 4. DISCUSSION

Rising global temperatures have an adverse effect on the production of major staple crops (Agnolucci and De Lipsis 2020; Jägermeyr et al. 2021) by hindering the development of seeds, as we observed in *Brassica napus* (Mácová et al. 2022). Understanding how plants perceive heat stress during seed development is important for breeding temperature-resilient crops. In our previous study, the occurrence of temperature-induced *in vivo* sprouting was reported in three spring cultivars of *B. napus* (Mácová et al. 2022), which affected seed viability and oil quality. Prolonged growth of Topas at high temperatures resulted in half of the seeds per silique undergoing SCR, a phenomenon rarely observed under optimal conditions. In this study, we aimed to understand the process resulting in the high-temperature-induced SCR phenotype in *B. napus* Topas cv. during late seed development, with a focus on seed coat development, cell wall modifications and biomechanics.

The integration of signalling pathways and the physical interactions between the three tissues that comprise the seed, the seed coat, the endosperm, and the embryo, determine the final size, shape, and dormancy state of the mature seed (Li et al. 2019). The *Arabidopsis haiku2-2* (*iku2-2*) *zhoupi-4* (*zou-4*) double mutant exhibits reduced seed viability due to developing seeds undergoing seed coat rupture (Creff et al. 2015), a phenomenon analogous to the SCR phenotype observed in Topas seeds developed at HT. The *iku2* mutation leads to elevated endosperm pressure, triggering premature mechanosensitive stiffening of the seed coat. This is likely associated with the accumulation of demethylesterified pectin, ultimately resulting in smaller *iku2* seeds (Creff et al. 2023). Seeds of *zou-4* have a persistent endosperm at the torpedo stage, even as the embryo expands in size, resulting in large seeds. Seed coat rupture in *iku2-2 zou-4* seeds may be a consequence of early stiffening of the seed coat together with increased pressure from the developing embryo and a persistent endosperm. We hypothesise that a similar process occurs in *B. napus* cv. Topas seeds, whereby high temperature alters the mechanosensing-dependent interaction between the embryo and the seed coat. From the earliest stages onward, high temperature accelerated embryogenesis, resulting in an approximately four-day developmental advancement by the onset of seed maturation (Mácová et al., 2022; and this study). Although seed size at 20 DAP was not affected, the enlarged embryos occupied most of the space in the seed cavity, thereby increasing mechanical stress on the surrounding seed coat and generating tension between the cells involved in seed size regulation. In *Arabidopsis*, the endosperm plays a significant role in modulating seed coat properties and determining seed size. Notably, endosperm-derived pressure has been proposed as a key mechanical component influencing seed coat integrity (Creff et al. 2023). Owing to technical limitations in *B. napus* Topas seeds, we were unable to assess whether high temperature similarly affected endosperm development or its mechanical contribution. This represents an important limitation of our study, as the proposed model partly relies on interactions between embryo growth, seed coat properties, and internal pressure dynamics. Consequently, we cannot exclude the possibility that endosperm-driven forces also contribute to the SCR phenotype under HT conditions.

The extensibility of the seed coat cells is determined by cell wall composition and the interactions among its structural components, which together permit embryo expansion. Transcriptomic analysis of the seeds revealed temperature-dependent differential regulation of cell wall-related pathways, with enrichment of genes associated with cellulose, pectin and xyloglucan metabolism, secondary cell wall thickening, and pectin modifications. The primary cell wall consists of cellulose and hemicellulose, embedded in a pectin-rich matrix (Cosgrove 2024), whereas the secondary cell wall can be further stiffened by lignin deposition. Cell wall stretching and thinning enable plant cell growth, which depends on the mechanical strength and dynamic extensibility of the wall. Wall strength and integrity are actively maintained by synthesis and modification of cell wall components (Peaucelle et al. 2011; Cosgrove 2024). Histochemical staining revealed increased pectin and lignin accumulation in the palisade layer of high temperature-developed seed coats. Consistently, immunolocalization using pectin-specific antibodies, together with cell wall profiling of early maturing seed coat, further demonstrated alterations in overall pectin composition, particularly in the palisade layer. Because the degree and pattern of pectin methylesterification influence cell wall rheology, these changes are likely to affect mechanical properties. Pectin, particularly its homogalacturonan (HG) subunit, undergoes dynamic methylesterification and demethylesterification processes that regulate wall extensibility (Du et al. 2022; Wang et al. 2025). The increased abundance of blockwise demethylesterified HG in the palisade layer at 14 and 18 DAP suggests enhanced Ca^2+^-mediated cross-linking, leading to cell wall stiffening and restricted cell expansion. This stiffening may explain why temperature-accelerated embryo growth was not accompanied by a proportional increase in overall seed size.

Nanoindentation analyses of mature seeds provided further insight into the mechanical properties of the seed coat cell walls. Nanoindentation of mature seeds showed that HT-developed seeds have both a lower reduced modulus (*E*_*r*_) and lower hardness (*H*_*IT*_) than CT seeds, indicating that the mature HT seed coat is mechanically more compliant and softer as a composite structure. At first sight, this appears to contrast with our biochemical observations, in which HT seed coat accumulate demethylesterified pectins (Figs 4-6), modifications typically associated with stiffening of individual cell walls, which correlates with an earlier seed growth arrest. We interpret these findings as complementary rather than contradictory. The immunolabelling and cell wall profiling report the chemistry of specific seed coat cell walls (palisade and epidermal) during maturation, whereas nanoindentation reports the mechanical behaviour of the whole seed surface at a coarser scale. Local chemical stiffening of individual cell walls can coexist with a structurally weaker overall seed coat, because the HT seed coat is also markedly thinner (Fig. 3). Thus, although the cell walls stiffen chemically, the thinner and structurally compromised seed coat is mechanically weaker as a whole and less able to constrain the expanding embryo. This mechanical imbalance is consistent with SCR we observed in HT seeds. Consistently, physical restriction of seed growth was previously shown to trigger seed maturation by arresting embryonic growth in *B. napus* cv. Reston (Rolletschek et al. 2024). Similar tubing experiments demonstrated that restricting embryo growth while promoting formation of a thicker palisade layer rescued the SCR phenotype, further substantiate our hypothesis. These results highlight how high temperature perturbs several key factors of seed development: accelerating embryo growth, increasing levels of pectin demethylesterification, and altering seed coat thickness. Ultimately, these changes disrupt the biomechanical coordination between embryo growth and seed coat resistance, compromising seed coat integrity.

## ACKNOWLEDGEMENTS

This work was financially supported by the project TowArds Next GENeration Crops, reg. no. CZ.02.01.01/00/22_008/0004581 of the ERDF Programme Johannes Amos Comenius. We acknowledge the Bioinformatics (supported by the NCLG research infrastructure LM2023067 funded by MEYS CR), Cell Imaging (supported by MEYS CR LM2023050 Czech-BioImaging), and Plant Sciences Core Facilities of CEITEC Masaryk University for their support in obtaining the scientific data presented in this paper. This work has benefited from the support of IJPB’s Plant Observatory platforms Po-Chem. The IJPB benefits from the support of Saclay Plant Sciences SPS (ANR-17-EUR-0007). This work was also supported by a grant from the French ANR (MECHASEED, ANR-23-CE13-0009) to BL. The nanoindentation tests were supported by the project LM2023039, funded by the Ministry of Education, Youth and Sports of the Czech Republic.

## AUTHOR CONTRIBUTIONS

HSR secured the funding. UP and HSR conceived the project, designed the experiments, and interpreted the data. UP performed the seed phenotyping, histology, transcriptomics, and immunolabelling of the seed coat. IU and JH optimised the protocol for Technovit embedding of oilseed rape seed and prepared the slides for histology. AMM analysed the transcriptomic data. AC, UP, and BL performed the immunolabelling of the seed coat. AV performed the cell wall profiling. VB and UP optimised the nanoindentation of Topas seeds. UP and HSR drafted the manuscript. All authors have proofed and approved the manuscript.

## DATA AVAILABILITY

The raw data generated during the current study and analysed in this published article (and its Supporting Information file) are available in the Zenodo repository (DOI: 10.5281/zenodo.19231242). The dataset supporting the conclusions of this article is deposited to the NCBI repository (BioProject accession number PRJNA1188476, https://www.ncbi.nlm.nih.gov/bioproject/PRJNA1188476).

## SUPPORTING INFORMATION

Additional Supporting Information may be found online in the Supporting Information section at the end of the article.

**Fig. S1.** Representation of seed phenotyping strategy in Brassica napus (Topas).

**Fig. S2.** Principal component analysis using gene expression data from the samples tested for differences between samples and biological replicates.

**Fig. S3.** Upregulated GO terms (Log_2_FC≥1) related to cell cycle regulation with the transcriptional response of Topas seeds to HT.

**Fig. S4.** Histological sections of Topas seeds focused on the seed coat, stained with Safranin-O for lignin.

**Fig. S5.** Quantification of xyloglucans in the Topas seed coat.

**Fig. S6.** Silicone tube mediated space restricted growth of seeds.

**Fig. S7.** Representative images of the 3D-printed mould for immobilizing the seed during nanoindentation.

**Fig. S8.** Represents the number of shared downregulated DEGs.

**Fig. S9.** Represents the number of shared upregulated DEGs.

**Fig. S10.** Load-displacement curves from the nanoindentation.

**Table S1**. RNAseq sample description.

**Table S2**. Full dataset for the DEGs for all the comparisons.

**Table S3**. Misregulated GO terms associated with plant cell wall, cell cycle and developmental pathways.

