## Supplemental Figures for "Long-term high temperatures affect seed maturation and seed coat integrity in *Brassica napus*"

The following Supporting Information is available for this article:

**Fig. S1.** Representation of seed phenotyping strategy in *Brassica napus* (Topas).

**Fig. S2.** Principal component analysis using gene expression data from the samples tested for differences between samples and biological replicates.

**Fig. S3.** Upregulated GO terms ($Log_2FC \geq 1$) related to cell cycle regulation with the transcriptional response of Topas seeds to HT.

**Fig. S4.** Histological sections of Topas seeds focused on the seed coat, stained with Safranin-O for lignin.

**Fig. S5.** Quantification of xyloglucans in the Topas seed coat.

**Fig. S6.** Silicone tube mediated space restricted growth of seeds.

**Fig. S7.** Representative images of the 3D-printed mould for immobilizing the seed during nanoindentation.

**Fig. S8.** Represents the number of shared downregulated DEGs.

**Fig. S9.** Represents the number of shared upregulated DEGs.

**Fig. S10.** Load-displacement curves from the nanoindentation.

**Table S1.** RNAseq sample description.

**Table S2.** Full dataset for the DEGs for all the comparisons (Excel table).

**Table S3.** Misregulated GO terms associated with plant cell wall, cell cycle and developmental pathways (Excel table).

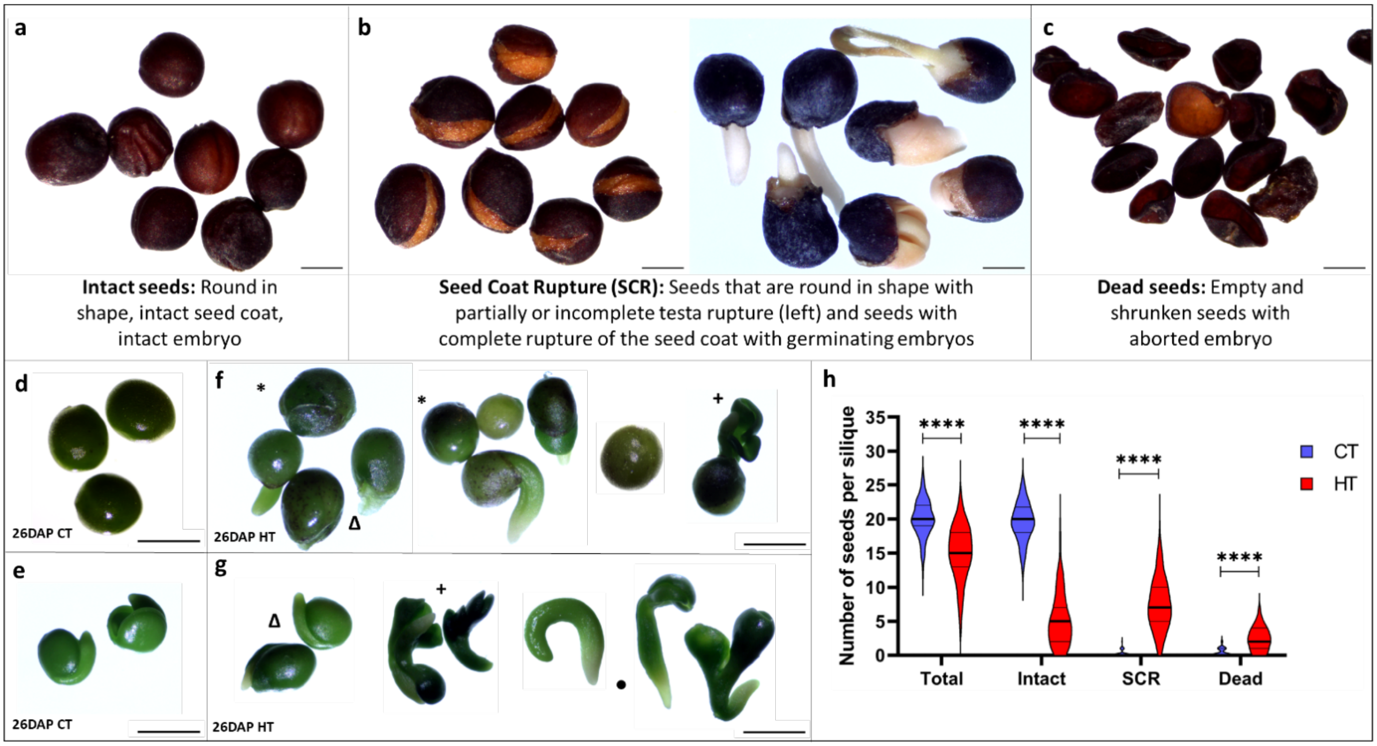


**Fig. S1.** Representation of seed phenotyping strategy in *Brassica napus* (Topas). (a-c) Representation of dry seed phenotyping strategy. (a) Intact seeds that are round in shape with an intact seed coat. (b) Seed Coat Rupture (SCR) phenotype with seeds that are round in shape with partially or incompletely ruptured seed coat (left) and seeds with complete rupture of the seed coat with germinating embryo (right). (c) Dead seeds, which are empty and shrunken seeds without an embryo. (d-g) SCR phenotype observed in Topas seeds before harvest. (d, e) 26 DAP seeds (d) and their isolated embryos (e) grown under CT. (f, g) 26 DAP seeds (f) and their isolated embryos (g) grown under HT. SCR was observed (*, Δ, +, ●). * Indicates seeds with seed coat rupture with intact embryo, Δ indicates seed coat rupture with radicle protrusion, + indicates seed coat rupture with cotyledon protrusion, and ● indicates embryos with abnormal development. (h) Quantification of the number of dry seeds per silique from CT (blue) and HT (red), showing the number of total, intact, SCR and dead seeds. Asterisks (*) indicate statistically significant differences in HT (**** corresponds to *p*-values of $p < 0.0005$).

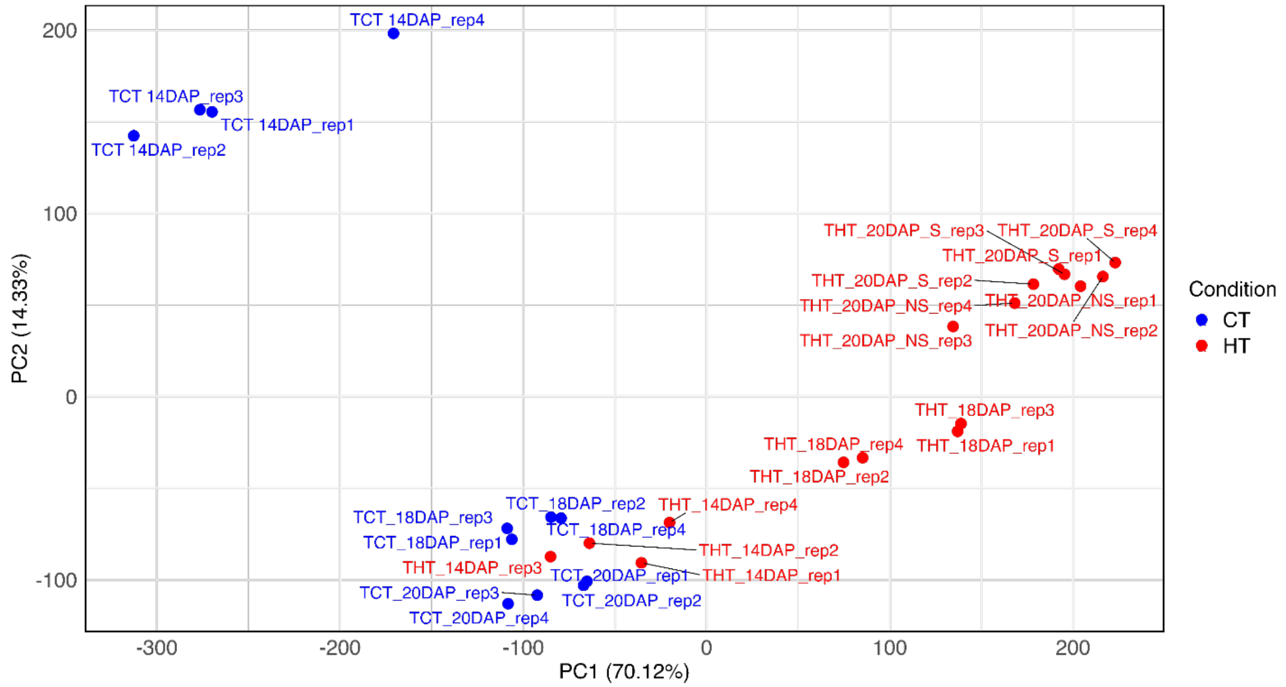


**Fig. S2.** Principal component analysis using gene expression data from the samples tested for differences between samples and biological replicates.

Principal Component Analysis (PCA) was performed in R on size-factor-normalized, variance-stabilizing transformed (VST) expression profiles, as implemented in DESeq2. PC1 and PC2 explained 70.12% and 14.33% of the total variance, respectively.

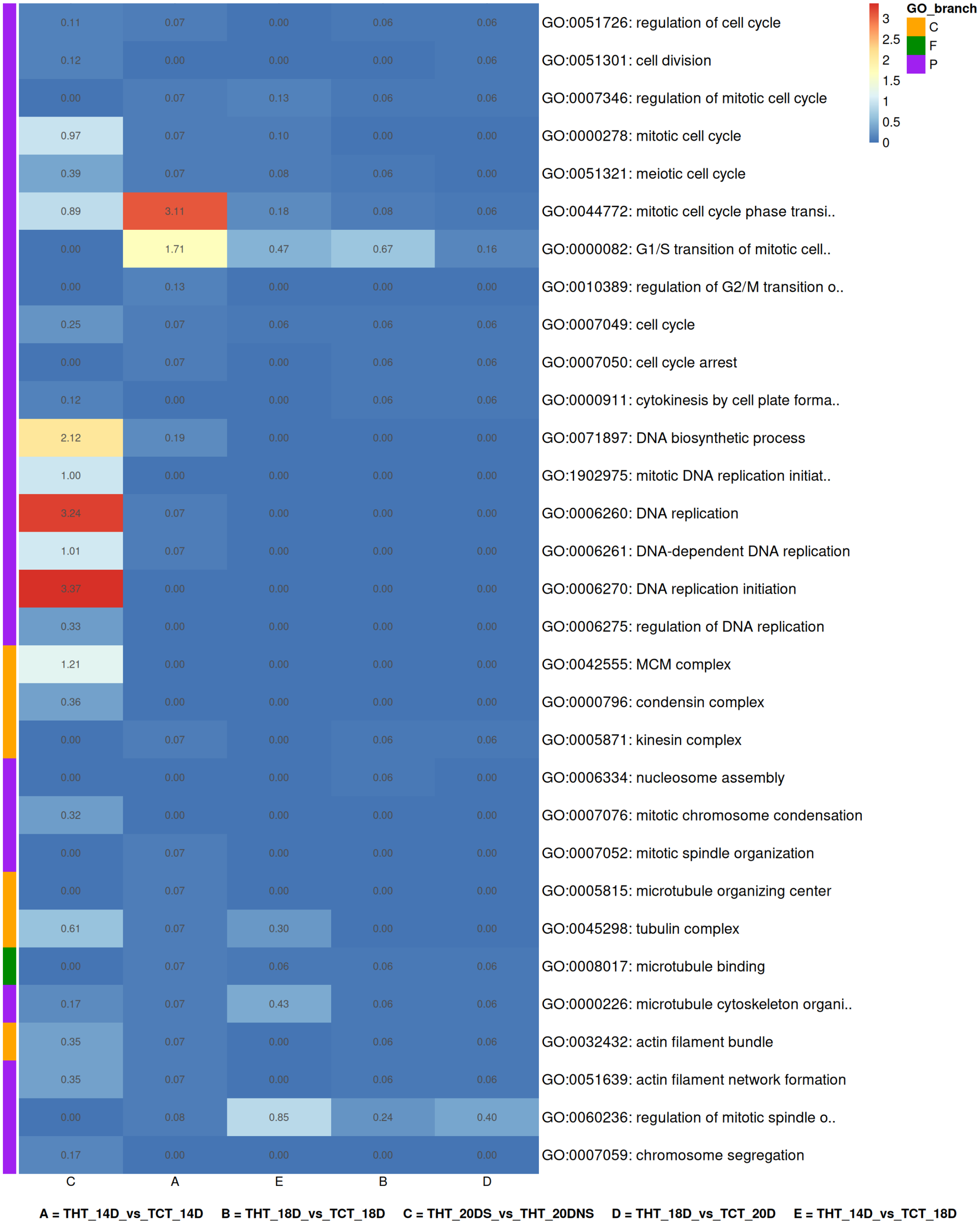


**Fig. S3. Fig. S3.** Upregulated GO terms ($Log_2FC \geq 1$) related to cell cycle regulation with the transcriptional response of Topas seeds to HT.

Values in the box depict $-Log_{10}$(q value).

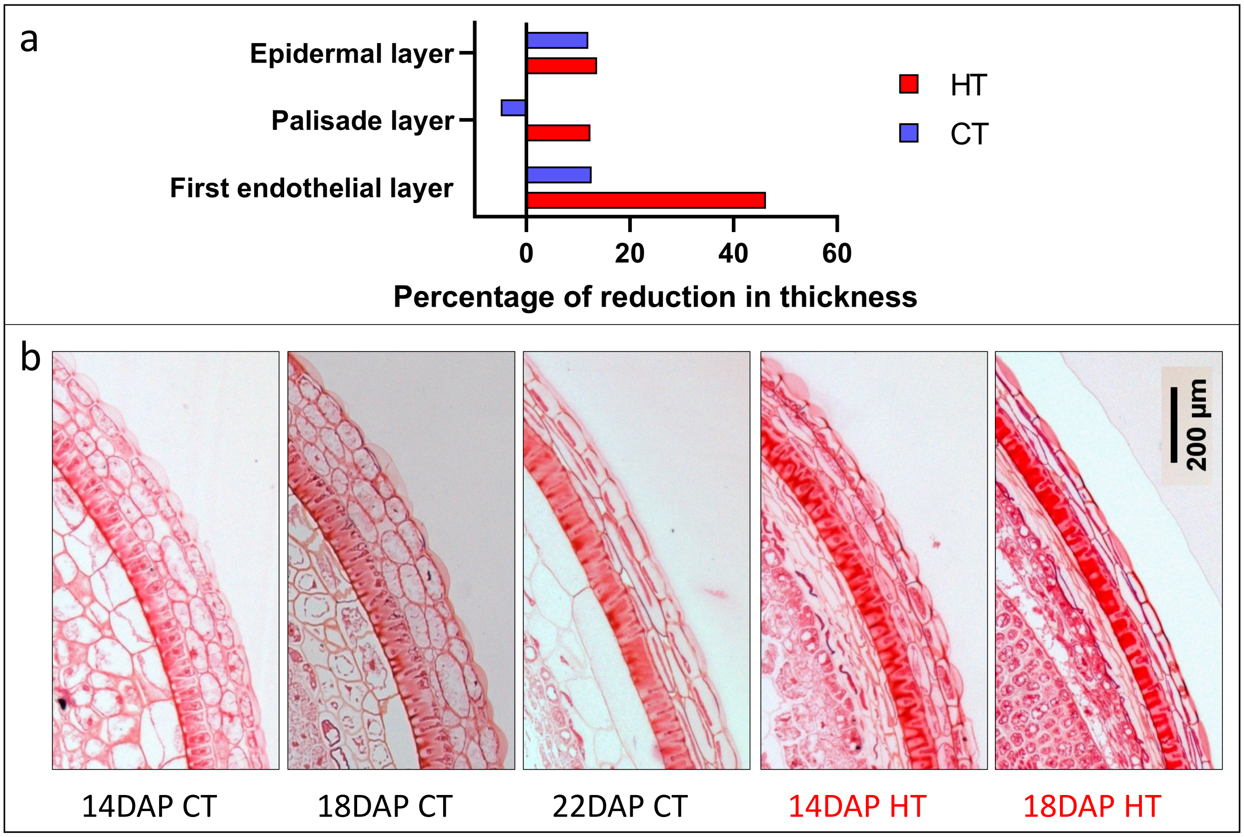


**Fig. S4.** Histological sections of Topas seeds focused on the seed coat, stained with Safranin-O for lignin. (a) Percentage of reduction in thickness as the seed aged from 14 to 20 DAP. Related to data presented in Figure 3. (b) Sections of 14, 18, and 22 DAP CT seeds, and 14 and 18 DAP HT seeds are shown. Scale bars represent 200 µm.

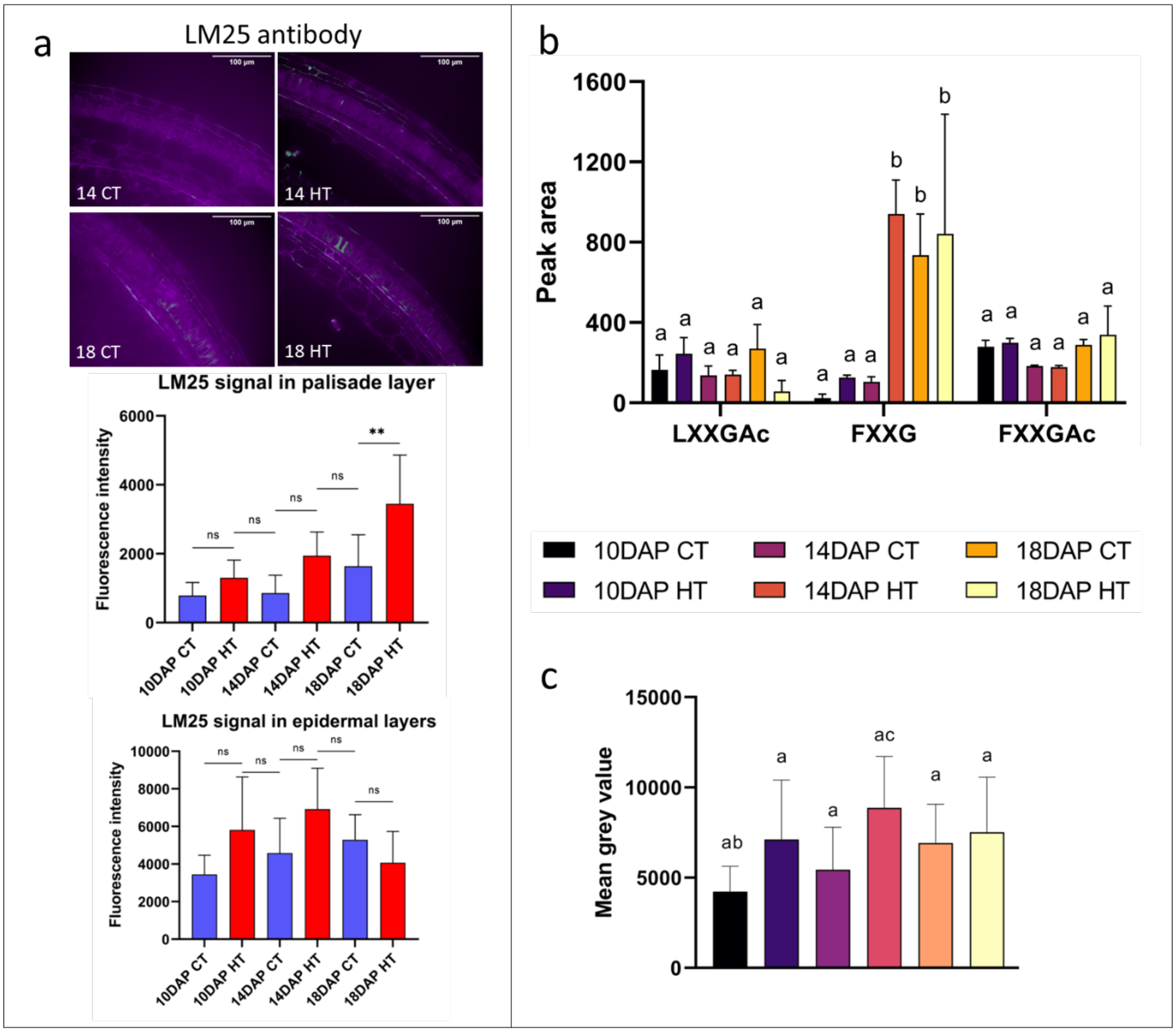


**Fig. S5.** Quantification of xyloglucans in the Topas seed coat. (a) Immunolocalization of xyloglucans using LM25 antibody on the seed coat of 14 and 18 DAP HT and CT seeds. The signal is visible in green, the cell structure is marked in magenta with Calcofluor White. Scale bars represent 100 µm. Fluorescence intensity was measured in epidermal and palisade cells in 10, 14 and 18 DAP HT and CT seeds. Asterisks indicate statistically significant differences in HT and between seed ages (*, **, *** and **** correspond to *p*-values of $0.05 > p > 0.01$, $0.01 > p > 0.001$, $0.001 > p > 0.0001$ and $p < 0.0001$, respectively); n.s., not significant. (b) Graph showing the measurements presented as peak area from the MS analysis for xyloglucans. (c) Graph showing the total intensity signal for xyloglucan from (a). Samples with the same letter do not significantly differ.

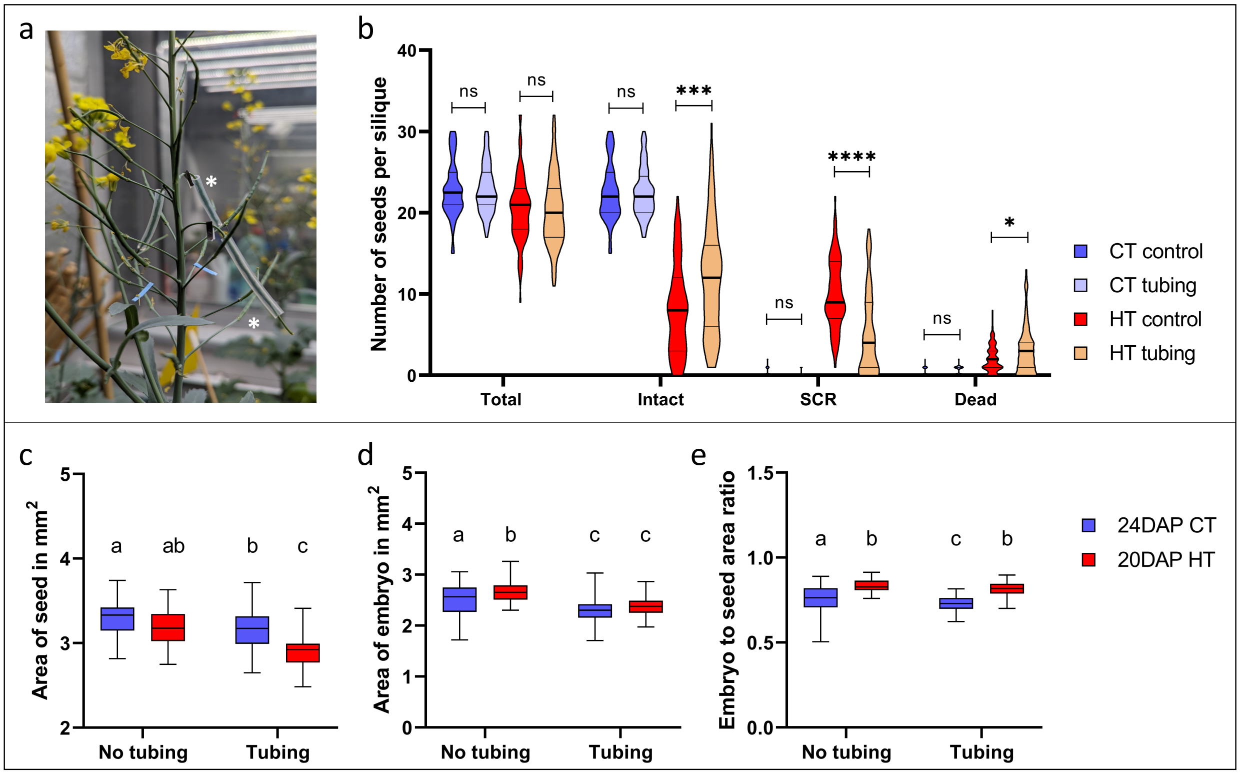


**Fig. S6.** Silicone tube mediated space restricted growth of seeds. (a) Representative image of a silique encased in a silicone tube. (b) A 3-mm silicone tube was applied to siliques from 12 DAP CT and 10 DAP HT. Dry seeds were quantified for the number of total, intact, SCR and dead seeds per silique. Asterisks indicate statistically significant differences in HT and between seed ages (*, **, *** and **** correspond to *p*-values of $0.05 > p > 0.01$, $0.01 > p > 0.001$, and $p < 0.001$, respectively); n.s., not significant. (c-e) Embryo and seed size measurements of Topas seeds from siliques encased in silicone tubes developed at CT and HT. (c, d) Seed (c) and embryo (d) area measurements in $mm^2$ of 24 and 20 DAP Topas seeds developed under CT (blue) and HT (red), respectively, to account for the same developmental stage. (e) Ratio of embryo-to-seed size in 24 DAP CT and 20 DAP HT seeds. Box-whisker plots show the distribution of area and ratio and the box indicates the interquartile range, the central line for median and the Tukey whiskers. Samples with the same letter do not statistically differ.

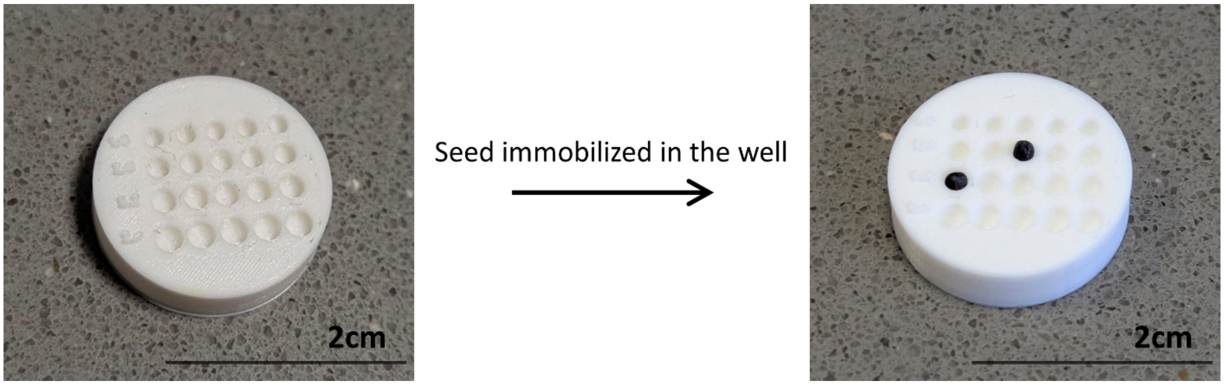


**Fig. S7** Representative images of the 3D-printed mould for immobilizing the seed during nanoindentation.

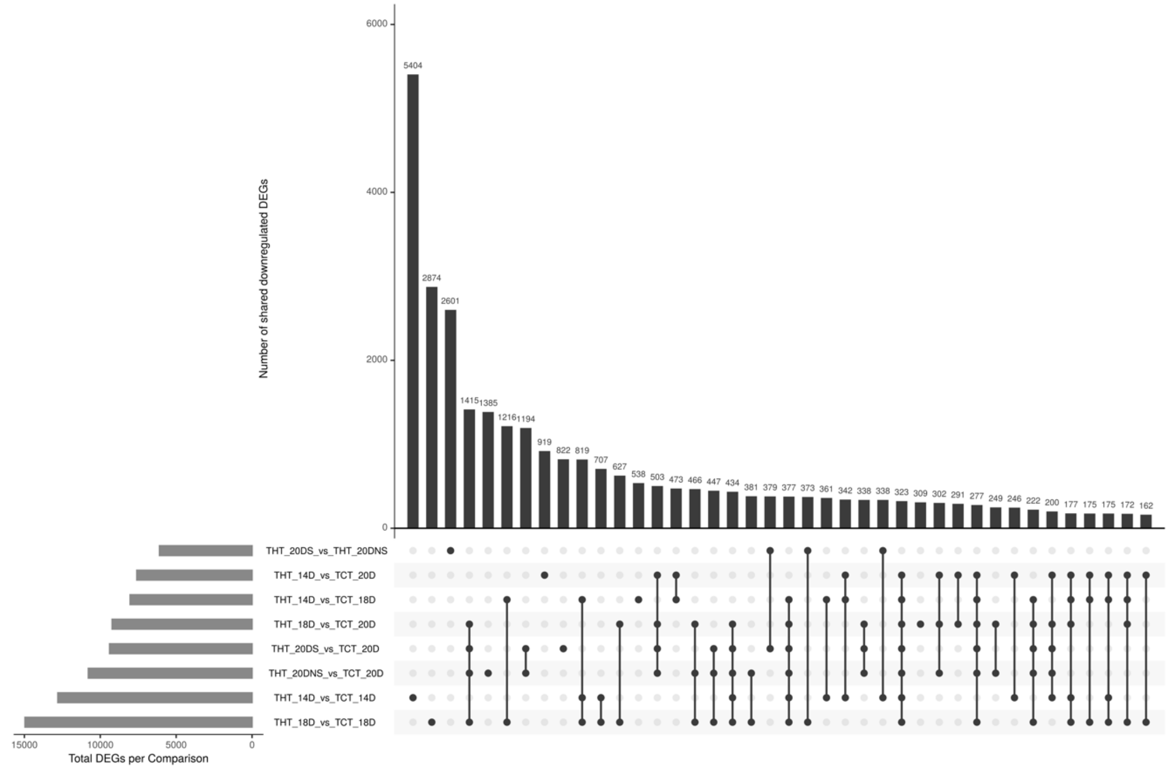


**Fig. S8** Represents the number of shared downregulated DEGs

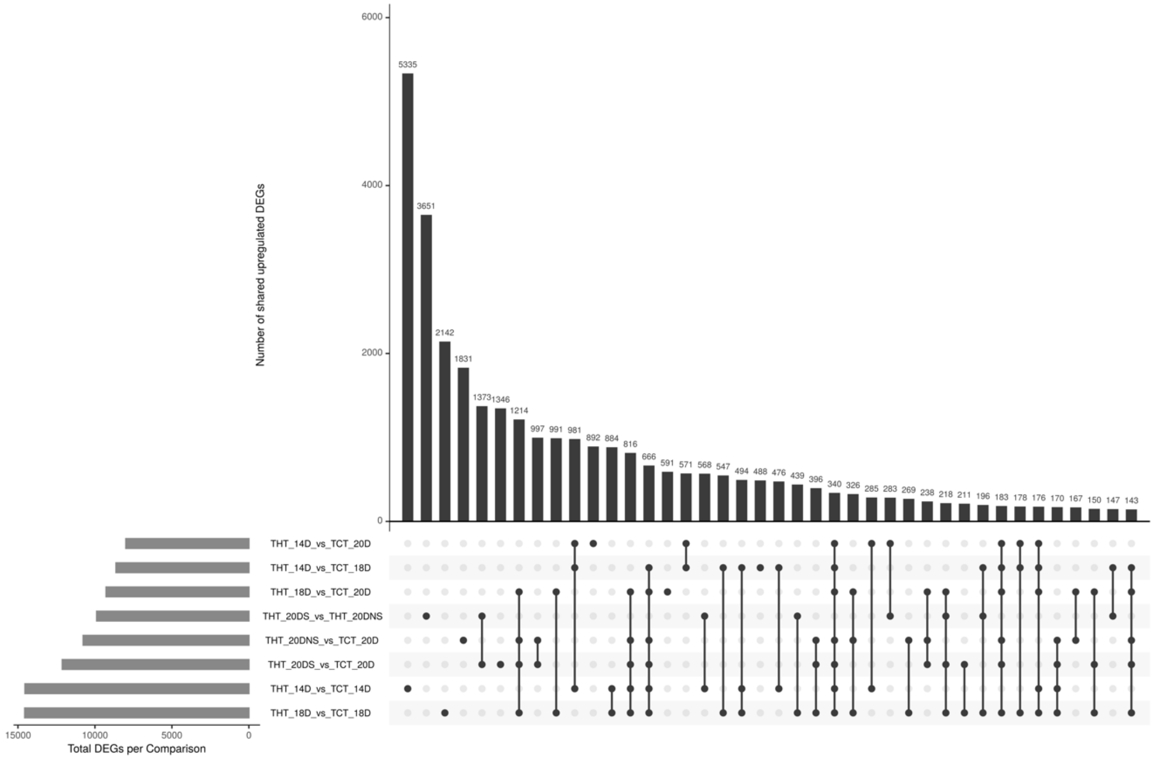


**Fig. S9** Represents the number of shared upregulated DEGs

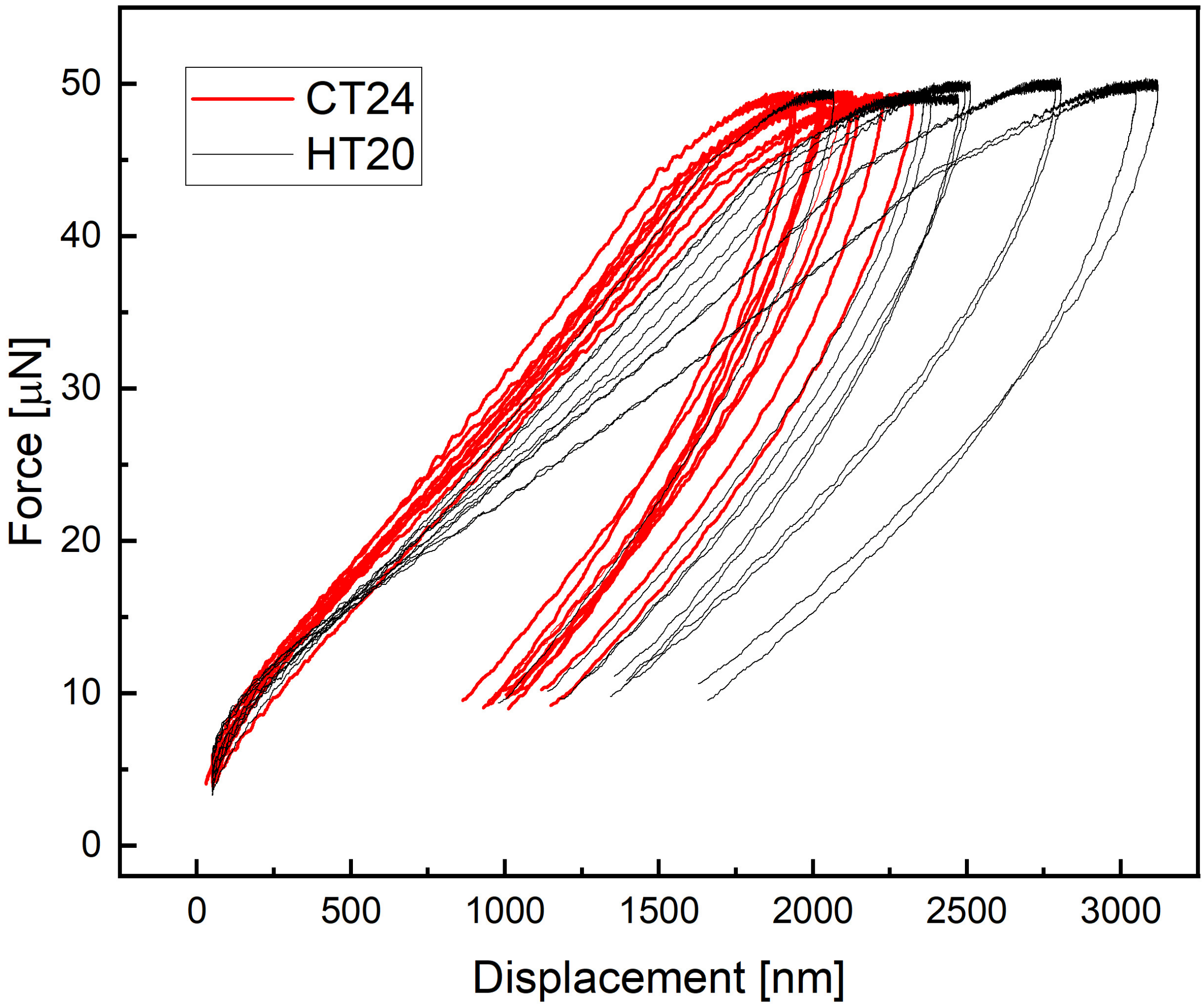


**Fig. S10** Load-displacement curves from the nanoindentation. Graph represents the comparison between the seeds of 24 DAP from CT (red) and 20 DAP from HT (black).

| Sample | condition | raw reads | preprocessed reads | uniquely mapped count | (%) | multimapped count | (%) |
|---|---|---|---|---|---|---|---|
| TCT_14D_1 | TCT_14D | 82,445,418 | 82,428,108 | 58,591,708 | 71.08 | 20,355,771 | 24.70 |
| TCT_14D_2 | TCT_14D | 92,209,285 | 92,189,952 | 65,503,046 | 71.05 | 22,782,105 | 24.71 |
| TCT_14D_3 | TCT_14D | 84,392,739 | 84,380,411 | 59,742,703 | 70.80 | 20,812,314 | 24.66 |
| TCT_14D_4 | TCT_14D | 63,807,782 | 63,799,082 | 45,336,600 | 71.06 | 15,692,511 | 24.60 |
| TCT_18D_1 | TCT_18D | 65,972,013 | 65,970,186 | 48,129,930 | 72.96 | 15,840,418 | 24.01 |
| TCT_18D_2 | TCT_18D | 63,004,795 | 62,995,974 | 45,397,632 | 72.06 | 15,062,201 | 23.91 |
| TCT_18D_3 | TCT_18D | 66,667,151 | 66,659,375 | 48,081,982 | 72.13 | 15,961,188 | 23.94 |
| TCT_18D_4 | TCT_18D | 62,660,193 | 62,652,931 | 45,051,127 | 71.91 | 14,966,731 | 23.89 |
| TCT_20D_1 | TCT_20D | 67,047,249 | 67,036,623 | 47,990,988 | 71.59 | 15,852,638 | 23.65 |
| TCT_20D_2 | TCT_20D | 67,249,609 | 67,239,841 | 48,331,955 | 71.88 | 15,985,039 | 23.77 |
| TCT_20D_3 | TCT_20D | 71,474,652 | 71,463,646 | 51,206,612 | 71.65 | 16,934,037 | 23.70 |
| TCT_20D_4 | TCT_20D | 74,579,007 | 74,567,670 | 53,470,636 | 71.71 | 17,714,329 | 23.76 |
| THT_14D_1 | THT_14D | 67,348,310 | 67,315,454 | 48,949,144 | 72.72 | 15,808,468 | 23.48 |
| THT_14D_2 | THT_14D | 68,610,483 | 68,595,024 | 49,170,149 | 71.68 | 16,277,066 | 23.73 |
| THT_14D_3 | THT_14D | 71,669,761 | 71,655,685 | 51,802,187 | 72.29 | 17,025,990 | 23.76 |
| THT_14D_4 | THT_14D | 62,316,943 | 62,297,640 | 44,469,960 | 71.38 | 14,720,636 | 23.63 |
| THT_18D_1 | THT_18D | 63,970,031 | 63,960,614 | 45,265,078 | 70.77 | 15,895,107 | 24.85 |
| THT_18D_2 | THT_18D | 74,578,172 | 74,562,728 | 52,358,723 | 70.22 | 18,796,498 | 25.21 |
| THT_18D_3 | THT_18D | 63,202,775 | 63,193,056 | 45,144,463 | 71.44 | 15,889,568 | 25.14 |
| THT_18D_4 | THT_18D | 72,666,475 | 72,656,752 | 50,886,635 | 70.04 | 18,234,971 | 25.10 |
| THT_20DNS_1 | THT_20DNS | 75,709,392 | 75,699,767 | 51,974,940 | 68.66 | 19,985,209 | 26.40 |
| THT_20DNS_2 | THT_20DNS | 73,069,275 | 73,060,722 | 50,213,631 | 68.73 | 19,330,352 | 26.46 |
| THT_20DNS_3 | THT_20DNS | 86,093,100 | 86,076,230 | 59,850,615 | 69.53 | 23,355,225 | 27.13 |
| THT_20DNS_4 | THT_20DNS | 81,883,774 | 81,863,352 | 55,079,025 | 67.28 | 22,334,329 | 27.28 |
| THT_20DS_1 | THT_20DS | 69,073,423 | 69,055,794 | 47,258,187 | 68.43 | 17,970,897 | 26.02 |
| THT_20DS_2 | THT_20DS | 69,782,647 | 69,770,711 | 47,514,805 | 68.10 | 18,358,543 | 26.31 |
| THT_20DS_3 | THT_20DS | 65,898,970 | 65,891,110 | 44,682,818 | 67.81 | 17,219,578 | 26.13 |
| THT_20DS_4 | THT_20DS | 63,818,858 | 63,809,808 | 43,102,515 | 67.55 | 16,646,496 | 26.09 |

**Table S1** RNAseq sample description

Condition assignment, number of raw and preprocessed reads, number and percentage of uniquely- and multi-mapped reads
